# Systematic humanization of the yeast cytoskeleton discerns functionally replaceable from divergent human genes

**DOI:** 10.1101/2019.12.16.878751

**Authors:** Riddhiman K. Garge, Jon M. Laurent, Aashiq H. Kachroo, Edward M. Marcotte

## Abstract

Many gene families have been expanded by gene duplications along the human lineage, relative to ancestral opisthokonts, but the extent to which the duplicated genes function similarly is understudied. Here, we focused on structural cytoskeletal genes involved in critical cellular processes including chromosome segregation, macromolecular transport, and cell shape maintenance. To determine functional redundancy and divergence of duplicated human genes, we systematically humanized the yeast actin, myosin, tubulin, and septin genes, testing ∼85% of human cytoskeletal genes across 7 gene families for their ability to complement a growth defect induced by deletion of the corresponding yeast ortholog. In 5 of 7 families—all but α-tubulin and light myosin, we found at least one human gene capable of complementing loss of the yeast gene. Despite rescuing growth defects, we observed differential abilities of human genes to rescue cell morphology, meiosis, and mating defects. By comparing phenotypes of humanized strains with deletion phenotypes of their interaction partners, we identify instances of human genes in the actin and septin families capable of carrying out essential functions, but apparently failing to interact with components of the yeast cytoskeleton, thus leading to abnormal cell morphologies. Overall, we show that duplicated human cytoskeletal genes appear to have diverged such that only a few human genes within each family are capable of replacing the essential roles of their yeast orthologs. The resulting yeast strains with humanized cytoskeletal components now provide surrogate platforms to characterize human genes in simplified eukaryotic contexts.

## Introduction

Gene duplication is regarded as one of the key drivers of evolution, contributing to the generation and accumulation of new genetic material within species^1, 2^. Duplication creates an initial multiplication of dosage and functional redundancy, but the trends dictating how duplicated genes retain function or diverge are still unclear. The processes governing the distribution of molecular roles within gene families also have important consequences for annotating genes, which generally takes advantage of sequence similarity and conservation over vast timescales of divergence to infer functions of homologous genes across species. Many individual studies have directly tested the conservation of function among orthologs from different species by swapping them from one species into another^3, 4^. However, only recently have efforts been made to test such functional equivalence more systematically, with several recent large-scale studies harnessing “the awesome power of yeast genetics” to systematically replace yeast genes by their human, plant, or even bacterial counterparts and assay for functional compatibility^5–10^. Although humans and yeast last shared a common ancestor nearly a billion years ago, these studies have demonstrated that substantial fractions (12-47%) of tested essential yeast genes could be replaced by their human equivalents^5–9, 11^. The ability of many human genes to functionally replace their yeast orthologs demonstrates the high degree of functional conservation in eukaryotic systems over billion year evolutionary timescales^9, 10^.

Previous humanization efforts in yeast have primarily focused on ortholog pairs with no obvious duplications within yeast and human lineages (1:1 orthologs), only partially testing the orthologs in expanded gene families^5–7, 9, 11^ and seldom beyond assaying impact on growth rate. In this study, to better understand functional conservation across expanded gene families in core eukaryotic processes, we focused on the major structural components of the eukaryotic cytoskeleton, including actins, myosins, septins, and tubulin genes. Genes constituting the eukaryotic cytoskeleton play key roles in critical cellular processes, mainly organizing the contents of the cell by dynamically controlling cell shape, positioning organelles, and transporting macromolecules including chromosomes across the cell through the generation of coordinated mechanical forces^12–14^. Importantly, cytoskeletal gene families have undergone large expansions along the human lineage, while being restricted to only a few family members in yeast (**Fig. 1**).

**Figure 1.**
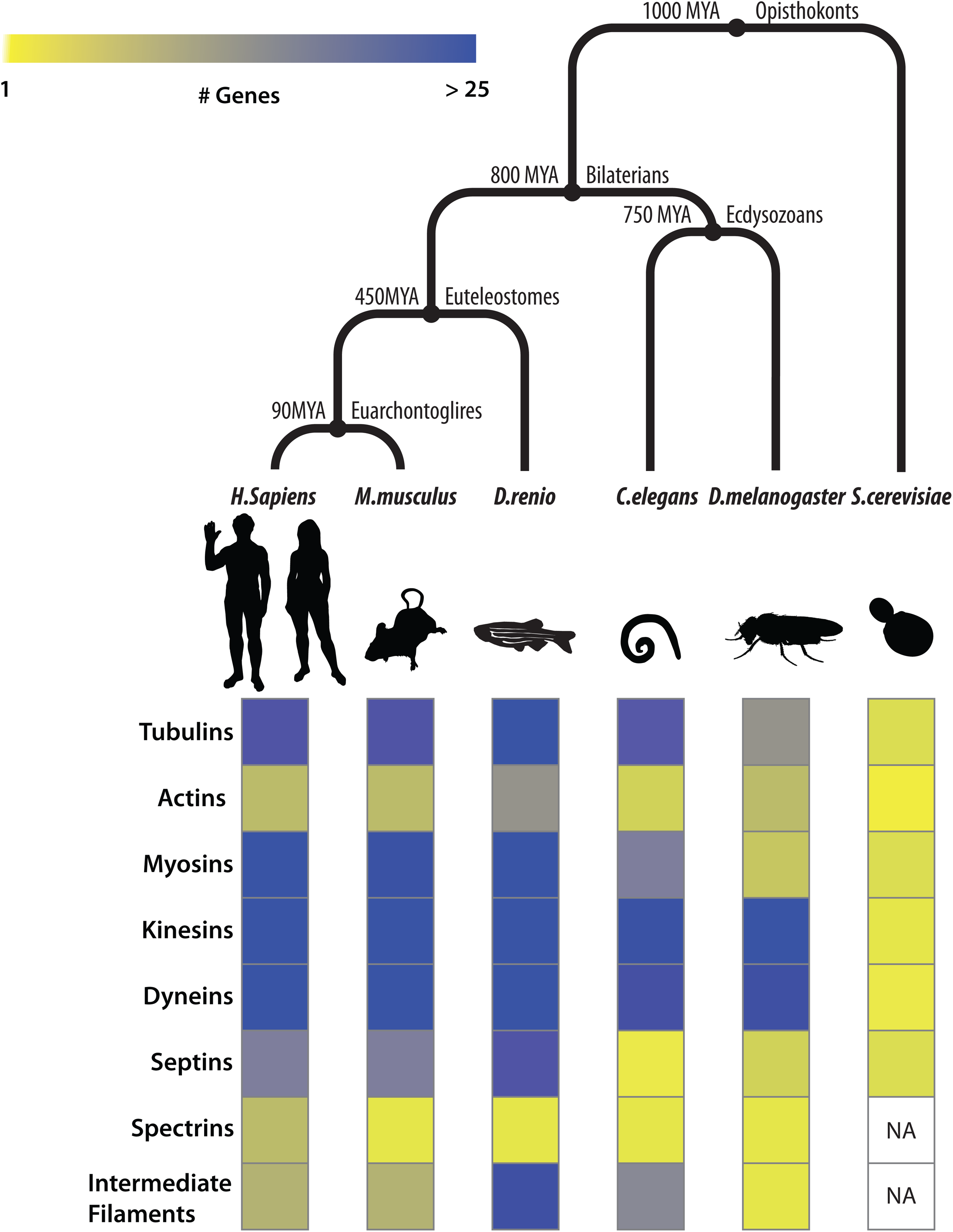
Orthologs in cytoskeletal gene families have undergone extensive duplications in Bilaterians. (A) (Top) Species divergence across opisthokonts. (Bottom) Heatmap depicting the number of orthologs in eukaryotic cytoskeleton gene families (rows) across species (columns). Cytoskeletal ortholog counts for model organisms curated from The Alliance of Genome Resources database^78^.

Advances in comparative genomics have shed light on the likely cytoskeletal components present in the last eukaryotic common ancestor (LECA)^12–14^. Additionally, phylogenetic profiling studies of the cytoskeleton have been highly informative in inferring gene loss and retention events across eukaryotic clades^13, 14^ (**Fig. 1**). Such studies suggest that the origins of the eukaryotic cytoskeleton predate eukaryogenesis and ancestrally trace back to primitive tubulin- and actin-like homologs in bacteria^13^. These components critical in cell division subsequently evolved to incorporate families of accessory motors and regulatory proteins expanding towards performing vital cellular roles, including phagocytosis, motility and vesicular transport, still evident across vast eukaryotic clades of life^13, 15–18^.

Though the cellular roles of human cytoskeletal gene families have been broadly elucidated, aided by studies in simpler eukaryotes^19–23^, the specific functions of their constituent family members in humans have to date still only been partially characterized^24–30^. Functional assays in human cell lines pose the challenge of functional redundancy, with buffering by other paralogs complicating the determination of paralog-specific roles within cytoskeletal gene families. The high degree of sequence conservation among paralogs within each cytoskeletal family make functional analysis of individual cytoskeletal genes directly in human cells both experimentally and computationally cumbersome. However, cross-species gene swaps have the potential to provide direct assays of individual paralogs within these expanded gene families, thereby revealing the extent to which present day orthologous genes retain ancestral function.

To understand the extent to which human cytoskeletal genes in expanded orthogroups retain cross-species functional equivalence, we systematically humanized major elements of the yeast cytoskeleton. We tested ∼85% (50/59) of all human genes from actin, myosin, septin, and tubulin families, using a combination of classical yeast genetics and CRISPR-Cas9 mediated genome editing to assay the replaceability of essential cytoskeletal orthologs, as initially determined *via* simple growth rescue complementation assays. Overall, we show that (13/59) members from 5 of 7 tested gene families (actin, heavy myosin, septin, β- and ɣ-tubulin) can indeed execute essential roles of their yeast counterparts. Within each replaceable family we show that several present-day human orthologs still possess functional roles of their respective opisthokont ancestors compatible in a yeast cellular context. Additionally, we characterized cellular phenotypes beyond growth and observed differential abilities among complementing human cytoskeletal genes to carry out non-essential cytoskeletal roles, including those in cell morphology, sporulation, mating, meiosis, and cytokinesis. Besides revealing human cytoskeletal orthologs capable of performing their core eukaryotic roles, yeast strains with human cytoskeleton components additionally serve as cellular reagents to study the specific roles of expanded cytoskeletal gene family members in a simplified unicellular eukaryotic context.

## Results

### Human cytoskeletal genes can functionally replace their corresponding yeast orthologs

Of the 324 cytoskeleton genes in yeast, 101 are essential for growth in standard laboratory conditions and possess identifiable human orthologs as determined by EggNOG^31^ and InParanoid^32^ (**Fig. S1**). We focused on orthologs constituting major structural elements of the eukaryotic cytoskeleton, including actin, myosin, septin, and tubulin gene families identifying 106 testable human-yeast ortholog pairs. By restricting our tests to elements of the yeast cytoskeleton essential for cell growth in standard lab conditions, we could initially assay replaceability of human genes in yeast *via* simple growth rescue assays. Since human cytoskeletal gene families have undergone multiple duplication events, we systematically swapped each human gene within a family in place of its corresponding yeast ortholog to assay if present day human genes could still complement the lethal loss of their yeast ortholog(s).

Complementation assays in yeast were carried out in 3 different ways (**Fig. 2**), wherein orthologous yeast genes could be (i) genetically segregated away after sporulation of a heterozygous diploid deletion strain^33–35^ (**Fig. S2A**) (ii) inactivated *via* a temperature-sensitive allele^22, 36^ (**Fig. S3A**) or (iii) endogenously replaced by homologous recombination repair with the human ortholog(s) following CRISPR-Cas9 cleavage ^8, 37^ (**Fig. S4A**).

**Figure 2.**
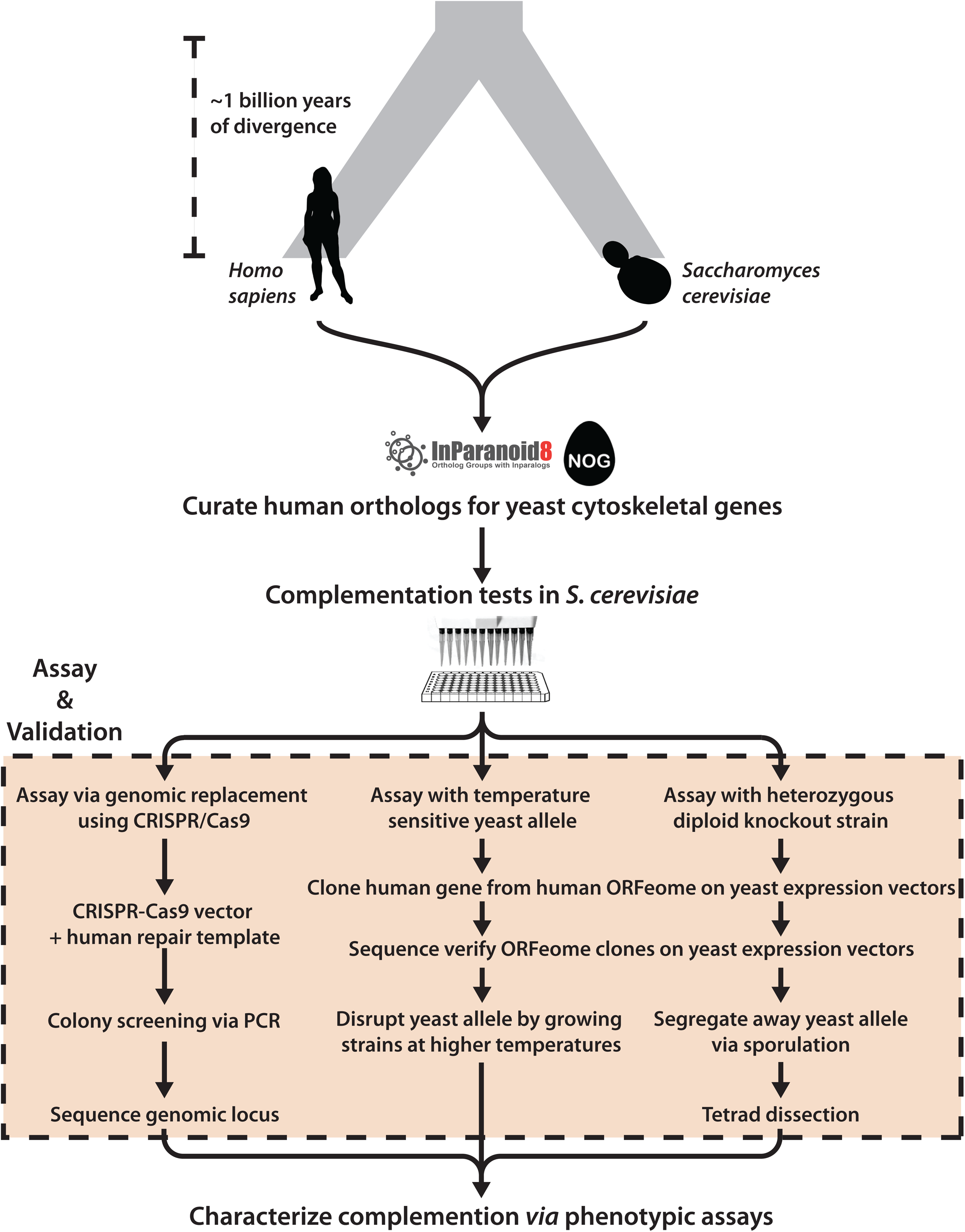
Overview of humanization assays. For each human-yeast ortholog pair (curated from inParanoid and EggNOG), complementation assays in *Saccharomyces cerevisiae* were performed using 3 strategies: (i) genomic replacement at the native yeast loci via CRISPR-Cas9, (ii) temperature-sensitive inactivation of the yeast allele, and (iii) sporulation of a heterozygous diploid deletion strain followed by tetrad dissection. Complementing human orthologs were further characterized using various phenotypic assays, including quantitative growth measurements, environmental stress tests, mating, and segregation assays.

Each assayed human gene was sequence verified and either sub-cloned into a single-copy (CEN) yeast expression vector transcriptionally controlled by a constitutive GPD promoter [in assays (i) and (ii)] or PCR amplified with flanking homology to the yeast locus of interest [in assay (iii)]. In cases involving heterozygous diploid sporulation assays, we leveraged the heterozygous diploid deletion collection^33^, wherein one copy of the yeast ortholog being queried has been replaced with a KanMX resistance module thus enabling selection of haploids with either the null (conferring G418 antibiotic resistance) or the wild-type allele (susceptible to G418) post sporulation (**Fig. S2B, C**). In cases of temperature-sensitive (ts) haploid strains, yeast genes of interest function optimally at the permissive temperature (25°C) but harbor mutations inactivating them at the restrictive temperature (37°C), allowing for temperature-dependent growth rescue complementation assays (**Fig. S3B, C**). Finally, in cases of endogenously chromosomal replacement, human genes were assayed by natively substituting their yeast ORF mediated *via* CRISPR-Cas9^37^ (**Fig. S4B, C**).

By taking advantage of full-length human cDNA clones from the human ORFeome^38–40^ and existing yeast strains with null (or conditionally null) alleles for the relevant orthologs^22, 22, 33, 34, 36^, we could successfully carry out 86 complementation assays across 89 human-yeast ortholog pairs, testing 50 out of 59 (∼85%) human genes across the 7 major cytoskeletal gene families. We observed that 13 (∼26%) of the 50 tested human genes could functionally replace their yeast orthologs in at least one of the three assay types, while 37 (∼72%) could not (**Fig. 2, Table S1, S2**). In particular, we found that 2 of 9 actin, 4 of 11 myosin (heavy chain), 4 of 13 septin, 2 of 9 β- and 1 of 2 ɣ-tubulin human genes complemented lethal growth defects caused by loss of their corresponding yeast orthologs, whereas genes from the human light chain myosin and α-tubulin families did not (**Fig. 3A**). Since the septin family had expansions in both human and yeast lineages, we systematically assayed all human septins against 4 yeast deletion backgrounds (*CDC3*, *CDC10*, *CDC11*, and *CDC12*), observing that complementing human septins functionally replaced only *CDC10*. Thus, within 5 of these 7 essential cytoskeletal gene families, at least one extant human gene could substitute for its yeast ortholog, indicating that the yeast and human genes both still executed the essential roles of their shared opisthokont ancestor.

**Figure 3.**
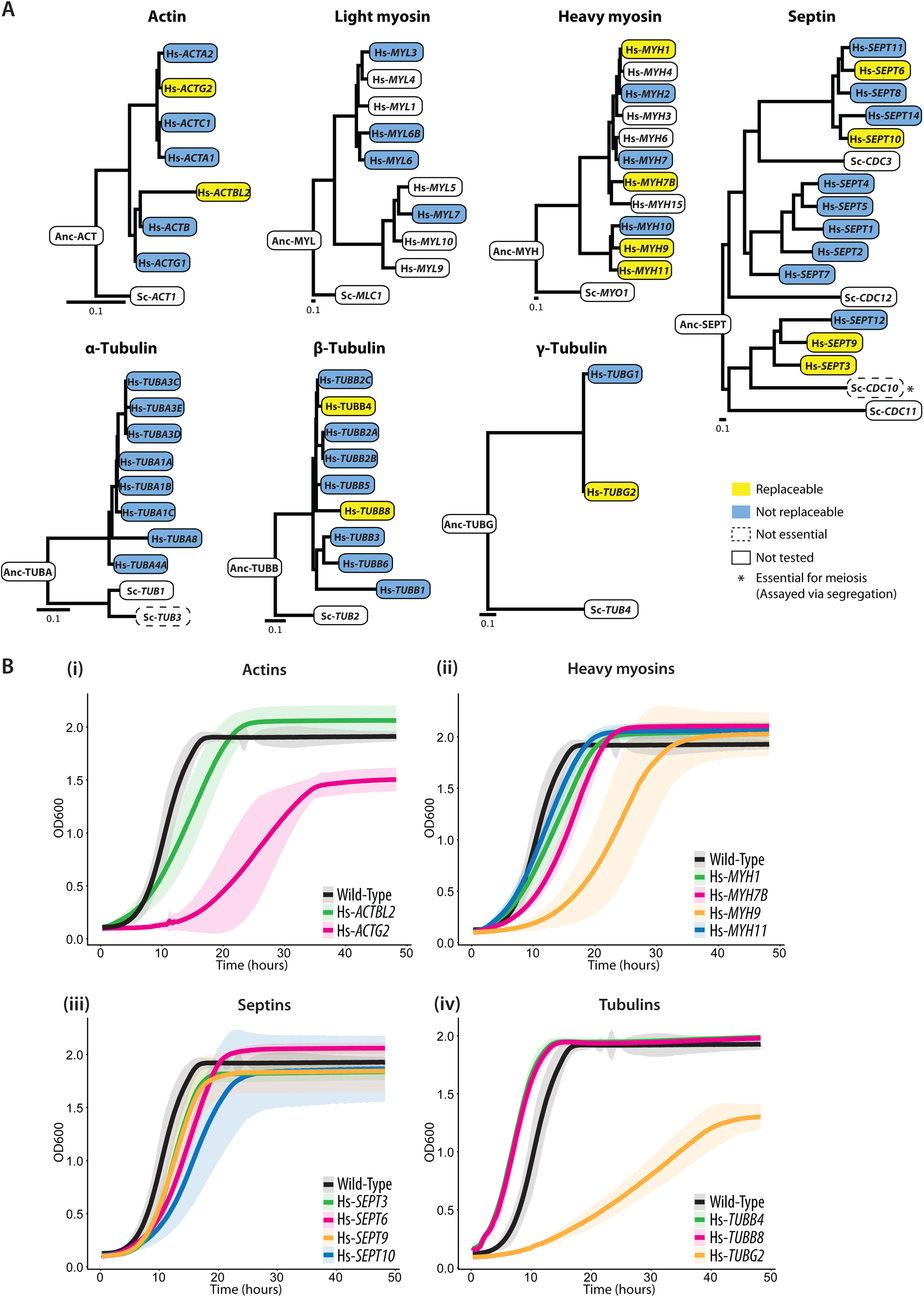
Human cytoskeletal genes replace their corresponding yeast orthologs. (A) 5 of 7 tested cytoskeletal families possess at least one functionally replaceable human ortholog. Neither of the tested human light myosin or α-tubulin genes could replace their corresponding yeast versions. Each human septin ortholog was tested for replaceability in 4 yeast septin null backgrounds (*CDC3*, *CDC10*, *CDC11*, *CDC12*), however, human septins complemented only *CDC10*, non-essential for mitotic growth but essential in segregation and mating. (Orthology relationships between human genes and yeast septins *SHS1*, *SPR3*, *SPR28* not shown). All phylogenetic gene trees were constructed with protein sequences of their respective orthologs (see **Materials and Methods**). Scale bars indicate expected substitutions per site. (B) Quantitative growth assays measuring absorbance at 600nm in (i) actin, (ii) heavy myosin, (iii) septin, and (iv) tubulin gene families. Except for Hs-*ACTG2*, Hs-*MYH9*, and Hs-*TUBG2*, all humanized yeast strains grow at similar rates in comparison to wild-type (BY4741). Since Hs-*ACTG2* and Hs-*TUBG2* were assayed via temperature sensitive suppression of the yeast ortholog, their inferior growth rates are likely due to a combination of inefficient replaceability and growth at higher temperatures of 37°C. Solid lines represent the mean and shaded boundaries demonstrate +/− standard deviation of 3 replicates each. For additional growth features see **Table S3**.

To assay robustness of complementation and better measure the extent to which human orthologs complemented in standard laboratory conditions, we performed quantitative growth assays on the humanized strains. We measured growth at 30°C (except for temperature-sensitive strains measured at 37°C) (**Fig. 3B**). While humanized septins showed no major growth defects, humanization of actin, heavy myosin, and tubulin genes led to minor growth differences. In particular, we found that replacement of yeast actin (*ACT1*) with Hs-*ACTG2* significantly slowed down growth and exhibited doubling times twice as long as wild-type yeast (**Fig. 3B (i), Table S3**), accompanied by reduced O.D. at stationary phase. Humanizing the yeast heavy myosin with Hs-*MYH9* did not affect growth to saturation phase with even its doubling time being twice the wild-type rate (**Fig. 3B (ii)**). In contrast, humanized ɣ-tubulin (Hs-*TUBG2*) strains showed drastically slow growth rates, doubling at one-quarter the rate of wild-type lab strains—and cells failed to reach comparable biomass at saturation (**Fig. 3B (iv), Table S3**). More broadly, we found that under standard laboratory growth conditions, most humanized strains exhibited growth profiles comparable to wild-type yeast.

To probe growth effects of humanization in more detail, we subjected the strains (except those assays performed in a temperature-sensitive strain background) to temperature stress, repeating the growth assays at low and high temperatures. While most humanized strains grew at wild-type rates at 25°C (**Fig. S9 bottom**), we observed differential effects at 37°C (**Fig. S9 top**). Specifically, yeast with human septins Hs-*SEPT6*, Hs-*SEPT9*, and Hs-*SEPT10* all reached stationary phase at a lower biomass compared to the wild-type yeast. Although septin genes complemented the lethal growth defect, they performed sub-optimally, especially under stress conditions, suggesting that they may be failing to complement other non-essential roles of the yeast orthologs, meriting a deeper examination of additional gene family specific phenotypes.

### Sporulation and meiotic roles of human β-tubulin are regulated by their genomic context in yeast

In yeast, β-tubulin (*TUB2*) is primarily associated with chromosome segregation during budding^21, 41, 42^ (asexual reproduction involving mitosis), mating^43, 44^ (sexual reproduction) and sporulation^45^ (starvation response involving meiosis). In our complementation assays, we observed that the human β-tubulins Hs-*TUBB4* and Hs-*TUBB8* only complemented the loss of *TUB2* when genomically inserted at the yeast *TUB2* locus but failed to do so in plasmid-based segregation assays in heterozygous diploid knockout strains. Notably, plasmid expression was controlled transcriptionally by a constitutive GPD promoter; both human genes failed to rescue its corresponding heterozygous diploid yeast gene deletion leading to sporulation defects producing either mis-segregated (**Fig. 4A - left panel**) or inviable spores (**Fig. 4B - left panel**). Therefore, replacing endogenous *TUB2* with human β-tubulins Hs-*TUBB4* and Hs-*TUBB8* at a minimum supported mitosis and asexual cell division *via* budding, but it was unclear if they supported meosis and sporulation.

**Figure 4.**
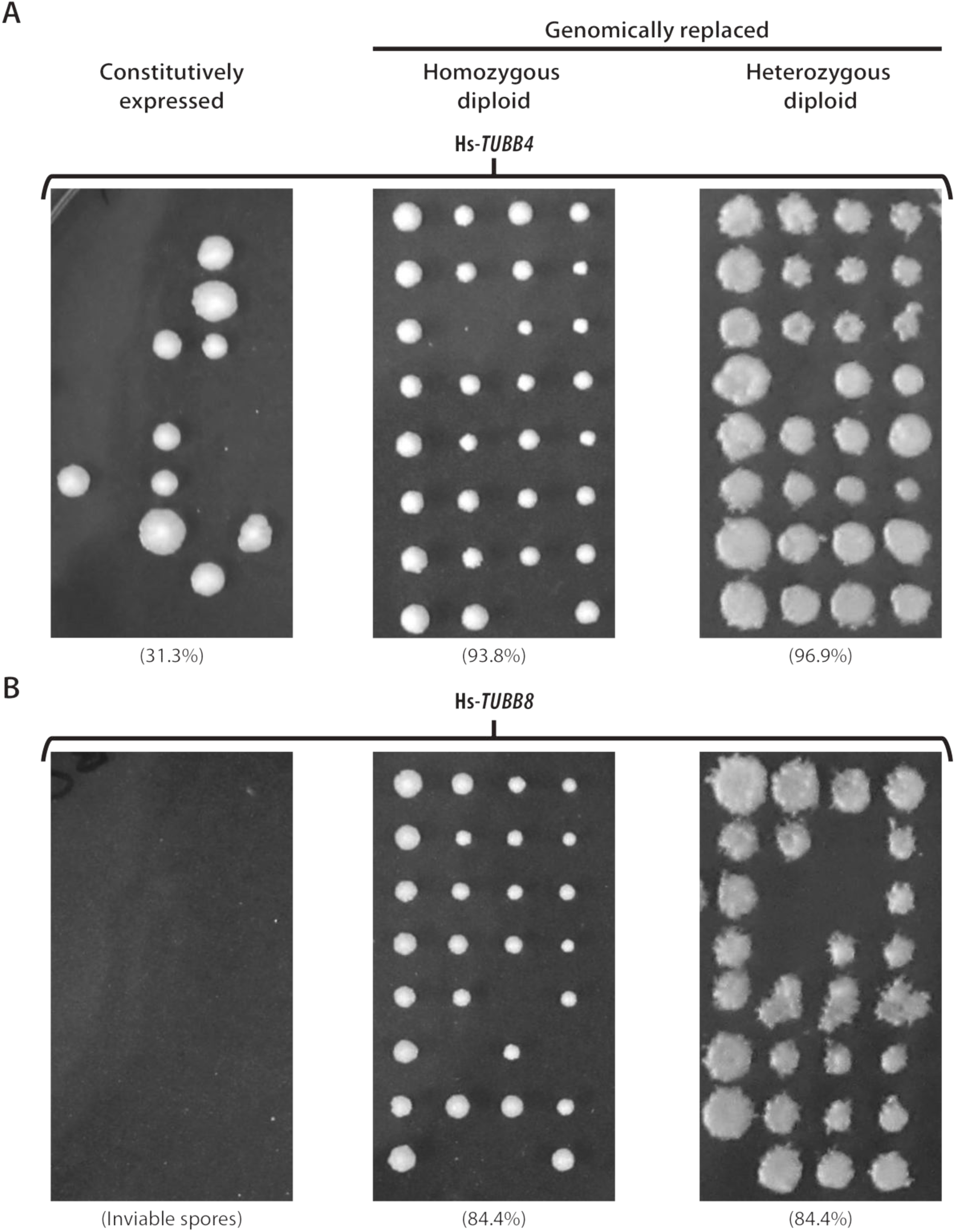
Replaceability of human β-tubulins Hs-*TUBB4* and Hs-*TUBB8* is determined by their native yeast expression/regulation. Efficiency of sporulation and segregation of both (A) Hs-*TUBB4* and (B) Hs-*TUBB8* tested in 3 different genetic backgrounds *via* tetrad dissection. Both constitutively expressed human β-tubulins Hs-*TUBB4* and Hs-*TUBB8* (controlled by a GPD promoter) fail to complement (left panel) whereas when replaced genomically could both grow and mate with wild-type yeast (middle panel). Diploids (both hetero- and homozygous) could also proceed through sporulation and meiosis similar to the wild-type yeast diploid strain (middle and right panel, see **Materials and Methods**). Spore viability indicated in brackets.

In order to determine whether replaceable human β-tubulins also rescued meiotic and sporulation specific roles of their yeast ortholog, we mated haploid humanized strains (harboring human genes at the corresponding yeast genes’ native genomic loci) with wild-type yeast. These humanized strains efficiently mated with wild-type yeast strains to produce viable diploids. We assayed if these heterozygously humanized strains could successfully perform meiosis by sporulating the diploids generated from our mating assays. We observed high spore viability for both Hs-*TUBB4* (∼96.9%) and Hs-*TUBB8* (∼84.4%) (**Fig. 4A & Fig. 4B - right panel**). Having shown that both replaceable β-tubulins can efficiently mate with the wild-type strains (yielding high sporulation efficiency similar to the wild-type diploids) (**Fig. S5A - right panel)**, we next asked whether strains homozygous for human β-tubulins could behave similarly. We sporulated the humanized homozygous β-tubulin diploid strains assaying for proper progression through meiosis *via* sporulation and found that these diploid humanized yeast strains also sporulated in both Hs-*TUBB4* (93.8%) and Hs-*TUBB8* (84.4%) cases. **(Fig. 4A & Fig. 4B - middle panel)** similar to their heterozygous (**Fig. 4A & 4B - right panels**) and wild-type diploids (∼87.5%) (**Fig. S5A - right panel** including 2:2 segregation of the MAT, LYS and MET loci after dissecting tetrads, see **Fig. S5B**). These results reveal that the genomic context and native regulation of β-tubulin are more critical to yeast than the species origin of the gene. The functional replacement of *TUB2* by two human orthologs, Hs-*TUBB4* and Hs-*TUBB8*, requires insertion into the native yeast locus, but nonetheless enables these genes to successfully perform their roles in meiosis and sporulation successfully, similar to the yeast β-tubulin *TUB2* (**Fig. S5)**

### Human septin orthologs and their isoforms can carry out *CDC10*’s meiotic and mating role

Next, we considered if human orthologs of yeast septins could also perform conditionally essential cytoskeletal roles in yeast. The yeast septin family is a particularly interesting case of gene family expansion as there have been duplications in both the human and yeast lineages, suggesting ancestral functions might have been distributed across paralogs in both species. Yeast possess 7 septin genes (*CDC3*, *CDC10*, *CDC11*, *CDC12*, *SPR3*, *SPR28*, and *SHS1*) while humans have 13 septin genes^46^ (**Fig. 1, 2A**). 3 of the 7 yeast septins (*CDC3*, *CDC11*, and *CDC12*) are essential for vegetative growth (budding) in standard lab conditions. On testing 39 of the 91 testable septin complementation pairs in each of the 3 septin null backgrounds, we found that none of the human orthologs assayed could complement *CDC3*, *CDC11*, and *CDC12*.

While the remaining 4 septins (*CDC10*, *SPR3*, *SPR28*, and *SHS1*) are not essential for vegetative growth, *CDC10* however, plays essential roles in meiotic cell division, mating, and spore health, with *cdc10Δ* strains exhibiting severe mating and sporulation defects^45, 47–49^. Indeed, we also observed that the diploid strain heterozygously null for *CDC10* sporulated poorly, almost always producing only 2 viable spores (**Fig. S2B(ii)**).

To identify human orthologs capable of rescuing sporulation defects caused by the loss of *CDC10*, we systematically assayed each of the 13 human septin orthologs. We observed that 4 of 13 septins (Hs-*SEPT3*, Hs-*SEPT6*, Hs-*SEPT9*, Hs-*SEPT10*) were capable of functionally complementing *CDC10*’s meiotic and sporulation roles (**Fig. 3**). Since *cdc10Δ* strains additionally show severe mating defects^47, 50^ (**Fig. S6A**), we next asked if the expression of individual human septin orthologs in a *cdc10Δ* strain could facilitate mating. We assayed this by cloning human septin genes into yeast expression vectors (**Materials and Methods**) and individually transformed *cdc10Δ* strains with each human septin clone. Subsequently, we mated each *MATa* transformant to a complimentary *MATα* strain (**Fig. 5A**). While only 4 of 13 tested human septins, Hs-*SEPT3*, Hs-*SEPT6*, Hs-*SEPT9*, and Hs-*SEPT10*, rescued *CDC10*’s essential meiotic role, we found that all assayed septins were capable of performing *CDC10’s* role in mating (**Fig. 5B**).

**Figure 5.**
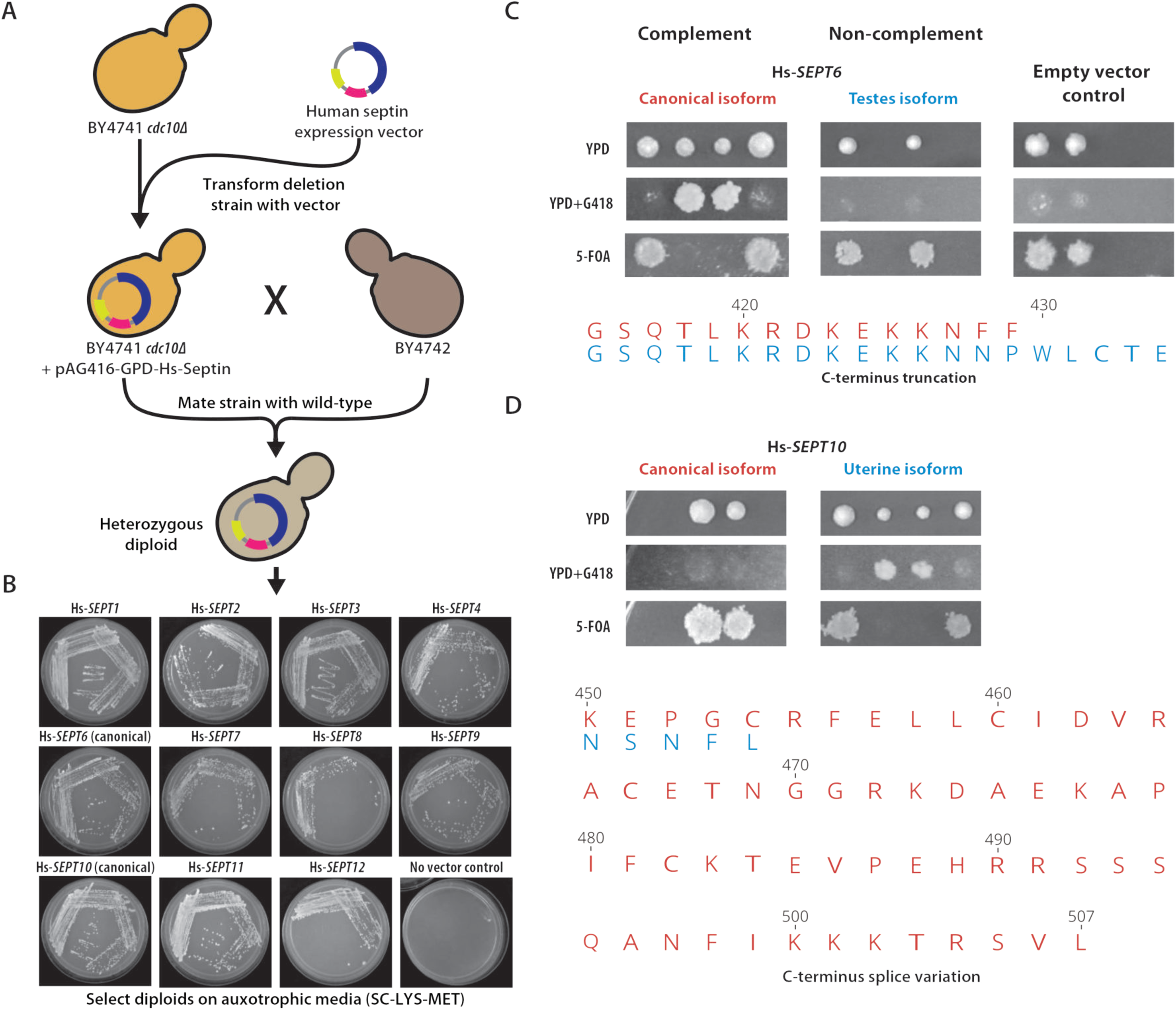
Human septins differentially rescue meiotic and mating roles of the yeast *CDC10*. (A) Mating rescue assay for BY4741 *cdc10Δ*. (B) All assayed human septins can efficiently mate with BY4742, whereas the empty vector containing BY4741 *cdc10Δ* fails to rescue the mating defect caused by deleting *CDC10*. (C) and (D) depict differential replaceability of human septin splice forms, Hs-*SEPT6* and Hs-*SEPT10*, respectively. The top panels demonstrate the heterozygous diploid deletion mutant segregation assay for the assayed isoforms while the bottom panels show the sequence alignment of the variation across the canonical (red) and tissue-specific (blue) isoforms tested for both Hs-*SEPT6* and Hs-*SEPT10*.

Unlike in yeast, human septins are known to exhibit a variety of different splice forms^16, 51^. Based on the availability of verified splice form (as per the Uniprot database) in the human ORFeome, we tested different isoforms of Hs-*SEPT6* and Hs-*SEPT10* observing contrasting results with both sets of isoforms. While the canonical isoform of Hs-*SEPT6* functionally complemented its yeast ortholog *CDC10*, its testes isoform with an extended C-terminus failed to do so (**Fig. 5C**). However, when testing the uterine and canonical isoforms of Hs-*SEPT10*, the opposite was true, with only the truncated uterine form functionally complementing *CDC10* (**Fig. 5D**). Thus for both, the presence of an extended C-terminus negatively affected the ability of the tested human septin isoforms to replace. Taken together, these results suggest that human septin orthologs can effectively rescue essential *CDC10* phenotypes in mating and meiosis.

### Humanized yeast strains differ in cell morphology

Eukaryotic cytoskeletal proteins dynamically control cell shape and morphology. We next asked whether humanizing elements of the yeast cytoskeleton would result in any visually obvious phenotypic changes to cell morphology. Using light and fluorescence microscopy, we imaged the 14 humanized strains generated and quantified cell shape and size across >22,000 individual cells (**Materials and Methods**). In the case of humanized heavy myosins, Hs-*MYH7B* and Hs-*MYH11*, we did not observe any obvious defects in the cell morphologies, but introducing Hs-*MYH1* and Hs-*MYH9* induced a slight size increase leading to more spherical cells (**Fig. 6A, S7A**). However, the effect, while significant, was small, and their median sizes were within 1% of wild-type (**Fig. 6A, S7B**). In contrast, strains with humanized actins, tubulins, and septins had visibly different cell morphologies (**Fig. 6A, S7**). Humanizing actin visibly reduced cell size, resulting in round/spherical cells with small buds (**Fig. 6A, S7A**). Complementing human ɣ-tubulin Hs-*TUBG2* showed the opposite phenotype, resulting in enlarged and ovoid cells (**Fig. 6A, S7**). The endogenously replaced human β-tubulin, Hs-*TUBB4*, also showed reduced cell size (**Fig. 6A, S7A**). Since α- and β-tubulins physically associate to form heterodimers we introduced human β-tubulins Hs-*TUBB4* and Hs-*TUBB8* into a strain expressing GFP-tagged α-tubulin^52^ (*TUB1-GFP*), allowing us to visualize microtubules and measure if chimeric yeast-human αβ-tubulin heterodimers assembled appropriately (**Fig. S8A**). Surprisingly, we observed visibly different cell morphologies when humanizing Hs-*TUBB4* (**Fig. S8A**), indicating a synthetic genetic interaction between the GFP tagged form of *TUB1* and Hs-*TUBB4*, absent from the interaction of Hs-*TUBB4* and native *TUB1*. Examining the distributions further, we observed a bimodal distribution of cell sizes (**Fig. S8B**) for Hs-*TUBB4* suggestive of cell cycle defects (as for *TUB2* Ser172 mutations^53^). In contrast, the cell morphology and microtubule assembly of the Hs-*TUBB8 TUB1-GFP* strain appeared similar to wild-type.

**Figure 6.**
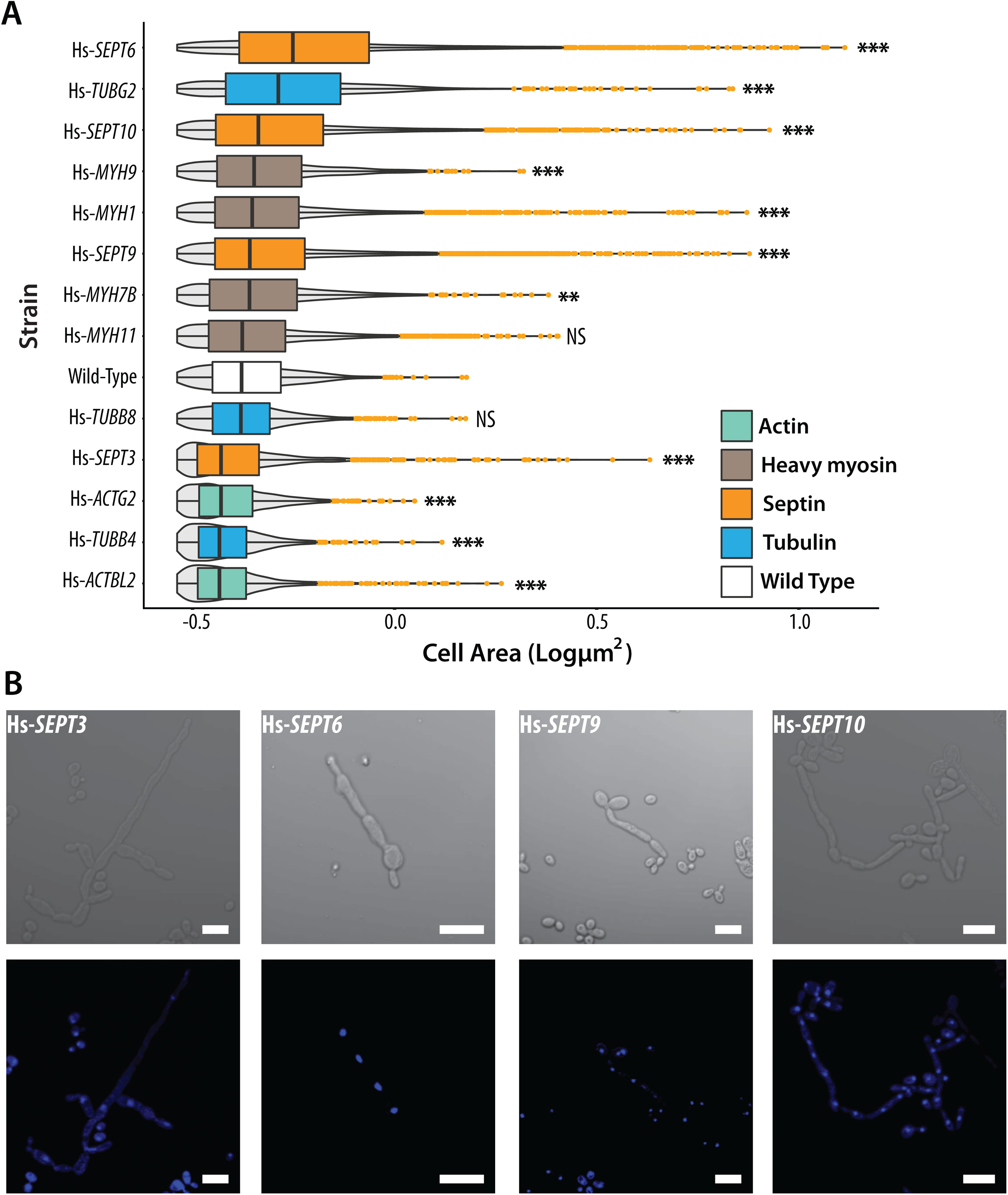
Yeast strains with humanized cytoskeletal components exhibit distinct cellular morphologies. (A) Humanized strains show varying cell sizes. Cell areas of humanized strains (in square pixels) are plotted on the X-axis. Grey violins indicate cell size distributions. Orange dots indicate outliers. Humanized actin strains show reduced cell size while myosins and tubulins (except ɣ-tubulin) largely remain unchanged. Significance determined by standard t-test with *** p≤10^−7^, **10^−4^≤p≤10^−3^, NS-Not significant). However, humanized septin strains show drastically elongated cellular morphologies. For additional bright-field images and significance of distribution tests see **Fig. S8.** (B) Magnified bright-field and DAPI-stained images of humanized septin strains exhibit elongated morphologies and are multinucleated as a consequence of defective cytokinesis. (Scale bars indicate 10 µm.)

All replaceable septin strains (Hs-*SEPT3*, Hs-*SEPT6*, Hs-*SEPT9*, and Hs-*SEPT10*) showed abnormal cell morphologies (**Fig. 6A, B, S7A**), notably reminiscent of elongated pseudohyphal forms often seen in pathogenic and invasive fungal species^54–56^. The fraction of elongated cells differed across human septins, with Hs-*SEPT3* producing lower proportions of elongated cells (**Fig. 6A, S7A**). To assay whether this effect arose from defective cytokinesis, we quantified the nuclei per cell by DAPI staining. We indeed observed that the cells were multinucleated (**Fig. 6C**). Despite this severe cell morphology defect, all replaceable human septins still enabled the strains to maintain growth rates comparable to wild-type strains (**Fig. 3D**). Taken together, we found that multiple human septins can rescue the essential meiotic and segregation roles of *CDC10*, but result in abnormal cell morphologies with delayed and/or defective cytokinesis.

### Humanized cytoskeletal orthologs phenocopy cell morphology defects observed by deleting interaction partners

In spite of rescuing lethal growth defects, the humanized strains showed visible cell morphology defects consistent with the known roles of the humanized genes. These defects suggested that the complementing human orthologs might be failing to perform some of the non-essential roles of the yeast genes for regulating cell shape and morphology. While the replaceable human cytoskeletal genes functionally complement the lethal growth defect caused by deletion of their yeast orthologs, we hypothesized that they failed to fully interface with their constituent interaction networks, thereby breaking key non-essential interactions regulating cell morphology. If true, this would imply that the humanization of a particular cytoskeletal yeast gene might phenocopy the deletion of its corresponding non-essential yeast interaction partner(s) (**Fig. 7A**).

**Figure 7.**
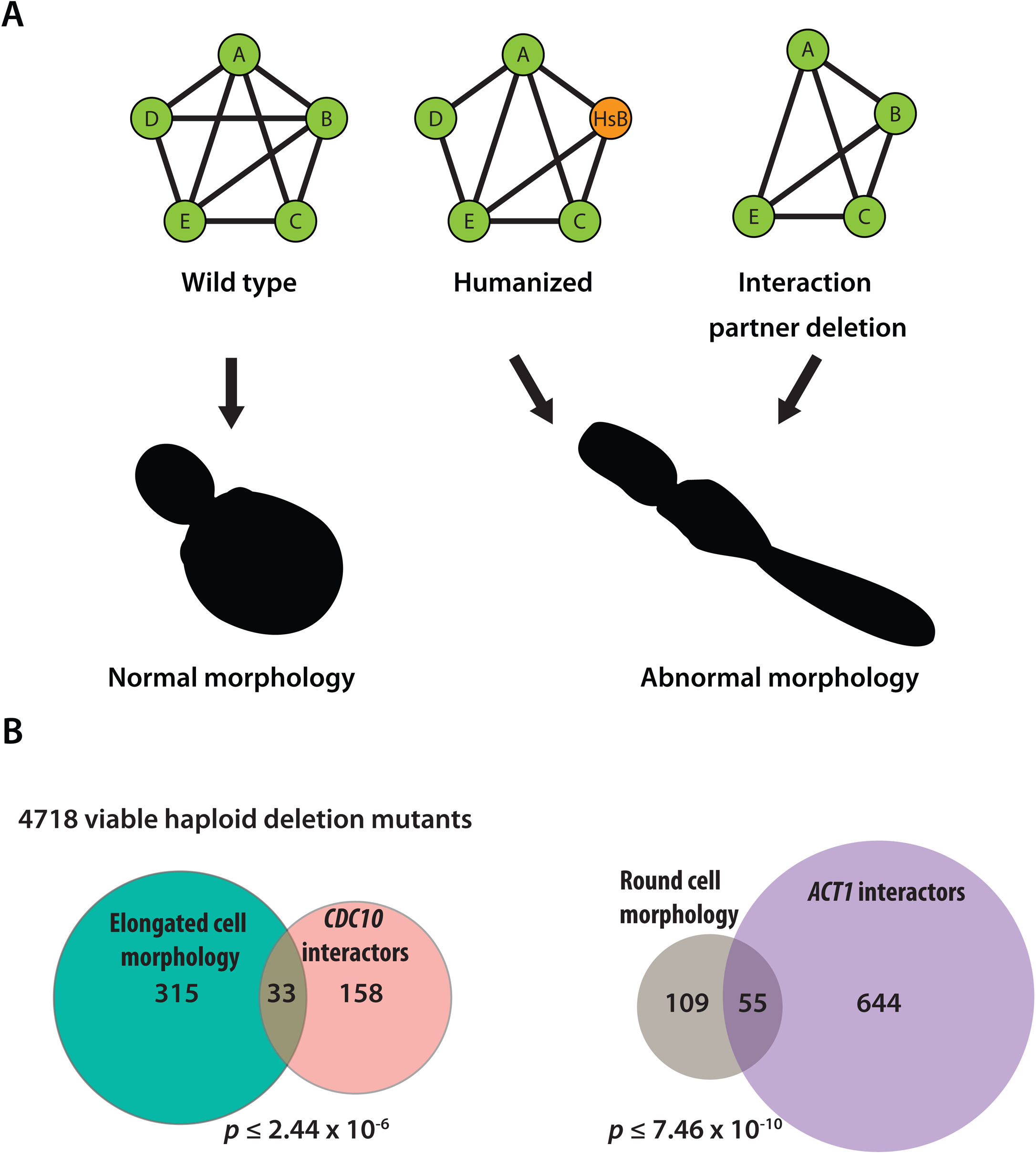
Humanized cytoskeletal yeast strains with abnormal cell morphologies phenocopy deletions of their corresponding yeast ortholog’s interaction partners. (A) Schematics of genetic interactions in the wild-type (Left), humanized (Middle), and interacting partner yeast deletion strains. Humanized yeast and yeast deletion strains are shown to cause similar abnormal cell shapes. (B) Cell shape parameters calculated from SCMD and genetic interactors of the replaced yeast ortholog curated from SGD show significant overlap (*p*-values determined by hypergeometric test).

To test this hypothesis, we curated genetic interaction partners of all humanizable yeast orthologs from the Saccharomyces Genome Database (SGD)^10, 55^ and mined the Saccharomyces Cerevisiae Morphological Database (SCMD)^56, 57^, a database cataloging the morphologies of ∼1.9 million cells across 4,718 haploid non-essential gene deletion backgrounds in yeast and measuring ∼ 501 cell shape parameters. Specifically, we considered the humanized strains showing drastic morphology changes and computed the ratio of long to short cell axes as a measure of cell roundness or elongation. Deletion strains with elongated phenotypes (*i.e.*, similar to humanized septin strains) would have higher axes ratios, and strains with round/spherical cells (*i.e.*, similar to humanized actin strains) would have axes ratios centered around 1. We found a significant overlap between the interactors of humanizable cytoskeletal genes and their observed phenotypes when deleted (p ≤ 2.44 × 10^−6^ for septin and ≤ 7.46 × 10^−10^, **Fig. 7B, Table S5**) suggesting that the replaceable human orthologs may indeed fail to maintain non-essential interactions controlling cell shape and morphology. It is possible that the amplification of cytoskeletal elements in the human lineage may have led to distribution of specific sub-functions and/or loss of interactions across duplicated human genes compared to the opisthokont ancestor. Our observations suggest that replaceable human cytoskeletal genes functionally complement the essential roles of their yeast ortholog, but simultaneously break key non-essential cytoskeletal associations, thereby phenocopying the deletion of their interaction partners.

## Discussion

Identifying how genes retain and/or distribute roles across their families is key to understanding the diversification of conserved genes across organisms. Cross-species gene swaps provide an opportunity to directly test the functional divergence of orthologous genes even over large evolutionary timescales. While *Saccharomyces cerevisiae* (Baker’s yeast) and *Homo sapiens* (humans) diverged from their opisthokont ancestor nearly a billion years ago, both species still share thousands of orthologous genes and high-throughput humanization assays in yeast have found that many human genes are capable of substituting for their yeast orthologs with rates up to 47%, depending on strains and assays^5–7, 9–11^. Most of these complementation tests until recently were performed in the absence of gene family expansions and revealed many humanizable systems, including the proteasome, and sterol and heme biosynthesis pathways^5–9, 11^. In this study, we sought to better understand the functional equivalence of human and yeast orthologs that play key structural roles in the eukaryotic cytoskeleton, particularly focusing on how gene family expansions in cytoskeletal lineages might have diversified in function across their respective gene families.

By systematically humanizing key structural components central to the yeast cytoskeleton, we determined that 5 of the 7 (∼71%) assayed eukaryotic cytoskeletal gene families could be successfully humanized by at least one human ortholog within a family. In all, 13 (26%) out of 50 tested human cytoskeletal proteins could at least partially substitute for the corresponding yeast gene and complement a lethal growth defect caused upon loss of the yeast ortholog (**Fig. 3A**).

Interestingly, these results are broadly consistent with our previous study, where 17 (∼28%) of 60 tested regulatory genes of the human cytoskeleton replaced their yeast orthologs^11^ (**Fig. S1**).

It is noteworthy that we were unable to successfully humanize yeast with any member of the myosin light chain or α-tubulin gene families, suggesting extensive functional divergence in yeast-human ortholog pairs resulting in failure to substitute for each other across species. This variation in replaceability was not merely explainable by obvious patterns across human genes (at least as captured by the variation in their expression patterns across tissues; **Fig. S10A**), nor was it explained by their degree of sequence similarity. As with previous yeast humanization studies^9^, sequence conservation among replaceable and non-replaceable human cytoskeletal genes did not significantly predict replaceability (**Fig. S10B**). While uncovering additional new properties predicting replaceability is beyond the scope of this study, future efforts at systematically constructing chimeric human/yeast genes have the potential to reveal which regions of the human/yeast orthologs are critical to maintain functional compatibility, perhaps enabling targeted humanization of specific domains and regions of genes.

With only slight differences in mitotic growth between wild-type and most humanized strains observed under standard laboratory growth conditions, our results suggest robust complementation of essential cellular roles in most cases. However, we subsequently found incomplete complementation of multiple non-essential cellular roles. We found that human β-tubulins Hs-*TUBB4* and Hs-*TUBB8* remarkably complemented *TUB2*’s roles even in sexual reproduction, including sporulation and mating. However, this was contingent on genomic integration under the native yeast regulation, consistent with previous studies showing that *TUB2* expression levels are tightly regulated^57^ with overexpression of β-tubulin leading to toxicity, chromosome loss and cell cycle arrest^58–60^.

For the septin gene family, which has expanded to consist of 7 yeast genes to 13 in humans, we found evidence for functional divergence across family members in both human and yeast lineages, with human genes tending to only fulfill a subset of specific roles of their yeast ortholog(s). While none of the human septin orthologs individually rescued essential roles of the yeast *CDC3*, *CDC11*, and *CDC12* genes, we found that 4 human septin orthologs (Hs-*SEPT3*, Hs-*SEPT6*, Hs-*SEPT9*, Hs-*SEPT10*) complemented *CDC10*’s essential roles in meiosis and sporulation. However, severe mating defects caused by deleting *CDC10* in a haploid strain background were completely rescued by every assayed member of the human septin family (**Fig. 5A, B**), indicating that all human septin orthologs executed the roles of their yeast counterparts to at least some extent. Extending this theme further, we saw that replaceability of Hs-*SEPT6* and Hs-*SEPT10* differed based on their protein isoforms, demonstrating functional divergence not just across human septin paralogs but also within splice forms of a gene (**Fig. 5C, D**). Taken together, such specific complementation patterns suggest a complex evolutionary trajectory and delegation of function across the septin gene family in eukaryotes. It remains to be seen if human septin orthologs can individually complement their non-essential yeast counterparts *SHS1*, *SPR3*, and *SPR28*. While septin gene family expansions have been understood to predominantly bring functional redundancy and robustness within its interactome, recent studies have identified tissue specific roles within human septin orthologs^61–63^. The mechanical roles facilitated by human and yeast septin gene families appear to be remarkably conserved albeit their involvement in seemingly unrelated processes across species^64^.

Over the course of these complementation assays, we observed diverse cell morphologies among humanized strains that were broadly gene family specific, with complementing myosins and β-tubulins largely remaining unchanged, but actin, ɣ-tubulin, septin families showing characteristic morphological differences as drastic as defective cytokinesis. Owing to the crucial role of the cytoskeleton in maintaining cell shape, we found that replaceable human cytoskeletal genes tend to perform its yeast ortholog’s essential roles equivalently while simultaneously breaking key non-essential cytoskeletal genetic interactions regulating cell morphology. In agreement with our findings, a recent study systematically determining cell size regulators in yeast found that 145 genes of ∼400 deletion strains were genetic interactors of actin^65^. Replaceable human septin orthologs, in particular, represent interesting and rather extreme cases of abnormal cell morphology. Previously, a study showed that introducing the *A. nidulans* septin *AspC* in *S. cerevisiae* induces a similar pseudohyphal morphology observed when substituting for *CDC12* in the septin ring^66^. More recently, a report^49^ demonstrated that doubly deleting *ELM1* and *FUS3* (both genetic interactors of *CDC10*) in yeast produces the same filamentous elongated morphology with similar cytokinesis defects observed when humanizing *CDC10*. In agreement with these studies, our results suggest that humanized cytoskeletal genes indeed phenocopy the deletion of their yeast ortholog’s interaction partners.

Systematic swaps of humanized cytoskeletal elements in yeast now provide a direct view of how compatible human orthologs likely are within their corresponding yeast interaction network(s), pointing to conserved and divergent interactions among eukaryotes. While our complementation assays tested the ability of human cytoskeletal orthologs to singly complement their yeast equivalents, combinatorial multi-gene swaps might enable humanization of entire yeast systems to study modularity and paralog-level cross-talk between different human cytoskeletal families in a genetically tractable eukaryote. Widening the scope of cytoskeletal humanization efforts to include accessory motors and chaperones, including kinesins and dyneins, could help advance our understanding of eukaryotic cytoskeletal evolution. It remains to be seen if constructing a fully human cytoskeleton in yeast would be feasible.

These humanized strains can now serve as cellular reagents to study complex human cytoskeletal processes in a simplified eukaryotic context, allowing functional roles of distinct family members to be assayed individually. Screening allelic variants and mutational libraries using these strains might enable the rapid identification of disease variants in a high-throughput manner^5–7, 9, 67, 68^, paving the way for a better understanding of the genetic and molecular basis of cytoskeletal disorders.

## Materials and Methods

### Curating human orthologs to yeast genes

Yeast genes to be tested for replaceability were curated from Saccharomyces Genome Database^4, 69^, and human orthologs were curated from the InParanoid^32^ and EggNOG^31^ databases both of which employ graph-based algorithms that recognize orthogroups between species by exhaustively performing an all-vs-all bidirectional BLAST search of all protein sequences and Hidden Markov Models to estimate ortholog groups respectively. We curated a total of 59 human cytoskeletal proteins with yeast orthologs in the 7 major cytoskeletal gene families (**Table S2**).

### Cloning human cytoskeletal ORFs

Human genes were extracted from the ORFeome^38^, a collection of *E. coli* strains, each containing a single human ORF cDNA cloned into a Gateway ‘entry’ vector^70^. Human genes cloned in this manner are flanked by attL recombination sites. To generate yeast expression vectors, the human entry vectors were isolated and subjected to Gateway LR reactions with a yeast Gateway ‘destination’ vectors followed by transformation into competent *E. coli* to obtain yeast expression clones. We used destination vectors from the Advanced Yeast Gateway kit^70^, specifically the pAG416-GPD-ccdB (CEN, URA) destination vector. Since the original version of this vector does not encode a stop codon immediately after the cloning region (which is also not encoded in ORFeome genes), this results in a ∼60 amino acid tail being translated to any protein expressed from it. To eliminate potential issues from the tail, we mutagenized the vector downstream of the cloning region to introduce a stop codon, thereby shortening the tail to six amino acids^9^. Prior to performing complementation assays, all expression clones were verified by Sanger sequencing to ensure there were no sequence errors or mutations in the human cytoskeletal genes prior to complementation assays in yeast.

### Assaying human cross-species complementation in yeast

#### 1. Tetrad dissection and analysis

In all, 40 human genes were assayed using tetrad dissection (**Fig. S2A**): The yeast heterozygous diploid deletion strain collection^33^ (obtained from ATCC) with one allele replaced with a Kanamycin-resistance (KanMX) cassette was used. Human genes in Gateway entry clones (pDON223) were curated from the Human ORFeome collection^38^ and cloned into yeast destination vectors. We transformed each human clone or an empty vector control into the appropriate yeast Magic Marker strain and grew them on SC–URA + G418 (200µg/ml) to select for the human clone (URA) and KanMX (G418) simultaneously. Transformants were then plated on GNA-rich pre-sporulation media containing G418 (200µg/ml). Individual colonies were inoculated into a liquid sporulation medium containing 0.1% potassium acetate, 0.005% zinc acetate, and were incubated at vigorous shaking at 25°C for 3-5 days. Following this, sporulation efficiency was estimated by microscopy, and successful sporulations were subjected to tetrad dissection and analysis using standard protocols^71^. Successful dissections were replica plated both on 5-FOA (for plasmid counter-selection) and YPD + G418 (for yeast null allele selection). Successful complementations consisted of 2:2 segregation with survival on YPD+G418 and failure to grow on 5-FOA (**Fig. S2B**). We subsequently performed quantitative growth assays (in triplicate) on the tetrads passing the 5-FOA and G418 segregation test (**Fig. S2B, S2C**). Each profile (**Fig. S2C**) was analyzed and quantified to detect any growth defects in yeast. 10 out of 40 human genes assayed in this manner functionally rescued yeast from the lethal phenotype. (**Table S2**).

#### 2. Yeast temperature-sensitive (ts) assays

Where available, temperature-sensitive (ts) strains from Li *et.al.*^22^ were used to assay human gene complementation. These strains grow ideally at lower permissive (22-26°C) but not restrictive temperatures (35-37°C). We transformed each human gene expression vectors and empty vector control plasmids into the corresponding ts strains (**Fig. S3A**). Transformants were plated on SC-URA media at both permissive and restrictive temperatures allowing us to control for transformation efficiency and expression toxicity while simultaneously testing for functional replacement by the human gene (**Fig. S3B**). Successful complementations consisted of transformants with the human gene surviving at both conditions (growth at permissive or restrictive temperatures) while those with the empty vector failing to survive at restrictive temperatures. Successfully humanizable strains were then subjected to quantitative liquid growth assays to monitor robustness of complementation (**Fig. S3C**).

#### 3. Genomic replacement *via* CRISPR-Cas9

29 human genes were assayed using a CRISPR-Cas9 mediated yeast genome editing protocol as previously described^8, 37^. For every yeast gene to be replaced endogenously, a minimum of 2 synthetic guide RNAs (sgRNA) with high ON-target and low OFF-target scores were designed using the Geneious v10.2.6 CRISPR-Cas9 tools suite. Selected sgRNAs were ordered as oligos from IDT and cloned into yeast CRISPR-Cas9 knockout plasmids from the yeast toolkit (YTK)^72^ to express a synthetic guide RNA, Cas9 nuclease and a selectable marker (URA) (see **Table S4** for guide sequence and primers). Wild-type yeast strains (s288c/BY4741) were transformed with a knockout plasmid and a repair template in the form of a PCR amplicon composed of the human open reading frame flanked with sequence homology to the targeted yeast locus (**Fig. S4A)**.

Transformants were selected on SC-URA medium to select for the knockout plasmid. While CRISPR-mediated double-stranded breaks are intrinsically lethal, targeted editing of an essential gene locus acts as an added layer of selection in replacing the human gene of interest, allowing cells to survive only if the human ortholog being assayed functionally replaces the yeast gene at the appropriate locus^8, 37^. Surviving colonies (**Fig. S4B**) obtained in the presence of repair templates were screened for successful humanization by colony PCR using primers outside the region of homology. Confirmed clones were Sanger sequenced and subsequently subjected to quantitative liquid growth assays to evaluate overall fitness. In the case of tubulins, all CRISPR-Cas9 assays were carried out in a BY4741 *tub3Δ* strain to avoid homologous repair of the *TUB1* locus by the *TUB3* gene. To generate diploid strains homozygous for human β-tubulin alleles, we replaced the yeast *TUB2* allele in our heterozygous diploids by re-transforming the CRISPR-Cas9 and sgRNA expression vector specifically targeting yeast β-tubulin allele (*TUB2*) and selected for viable strains without supplying an external repair template, forcing homology-directed repair of the *TUB2* lesion by the human ortholog(s) on the other homologous chromosome.

### Growth Assays

Liquid growth assays were performed in triplicate using a Biotek Synergy HT incubating spectrophotometer in 96-well format. All humanized strains were pre-cultured to saturation in YPD and diluted into 150µL of medium to finally have 0.05-0.1 × 10^7^ cells/ml. Each growth assay lasted for 48 hrs, with absorbance measured at 600nm every 15 min. For human genes assayed using heterozygous diploid deletion collections, growth assays were performed in YPD, SC-URA, and YPD+G418 medium to confirm retention of the plasmid and selection of the deletion allele respectively. For CRISPR assays, humanized strains were grown in YPD. Human genes assayed *via* temperature-sensitive alleles were grown at both temperatures (permissive and restrictive). Growth curve data was processed in Rstudio and plotted using the ggplot2 package.

### Microscopy and image analysis

For DAPI staining, cells were grown in liquid medium to saturation and fixed in a final concentration of 3.7% (v/v) formaldehyde for 10 minutes at room temperature. Cells were then washed with water and 1M sorbitol after which they were resuspended in sorbitol (1x of the original volume) and fixed in 50% ethanol (final concentration). The cells were washed and resuspended in 1M sorbitol. DAPI was then added to the solution to a final concentration of 1µg/ml and incubated for 5-10 min at room temperature.

Cells were imaged using a Nikon TE-2000-E inverted microscope with a Apo 40x/NA 0.95 objective and Cascade II 512 camera (Photometrics), Lambda LS Xenon light source and Lambda 10-3 filter wheel control (Sutter Instrument) with a motorized stage (Prior Scientific). All imaging and parameters were set via the Nikon NIS Elements Imaging Software. Images were captured at 1 frame per second through a 89000ET filter set (Chroma Technology) with channels “DIC L”, “FITC” (Ex 490/20, Em 525/36). GFP fluorescence images were collected with an exposure time of 1s. DAPI images for the humanized septin yeast strains were obtained using a Zeiss LSM 710 confocal microscope with a Plan-Apochromat 63x/1.4 oil-immersion objective with standard DAPI (Ex 350/50, Em 455/50) wavelength setting and operated using Zeiss ZEN Microscope software.

Cell size measurements were performed with a minimum of 10 fields of view and 2000 cells per strain. Image analysis and quantification was performed using FIJI/ImageJ^73, 74^. For quantifying cell size, edge detection scripts were written as Python scripts and FIJI macros.

### Mating assays

Since the genes assayed via CRISPR were tested in BY4741 (genotype MATa *his3Δ1 leu2Δ0 met15Δ0 ura3Δ0*), the humanized strains were mated to a BY4742 derivative (genotype MATα *his3Δ1 leu2Δ0 lys2Δ0 ura3Δ0 mkt1*-D30G *rme1*-ins308A tao3-E1493Q) so that diploids could be selected on SC-LYS-MET medium. Strains were first patched on a YPD plate. The humanized strain being assayed and its complementary mating strain were streaked perpendicular to each other and were grown overnight. These were then replica plated on SC-LYS-MET media to select for diploids. Diploids were grown overnight on GNA rich pre-sporulation medium. Individual colonies from this plate were inoculated into a liquid sporulation medium containing 0.1% potassium acetate, 0.005% Zinc acetate, and were incubated at vigorous shaking at 25°C for 3-5 days. Spores were dissected on a YPD plate using a similar protocol, as described previously. Dissected spores were replica plated on SC-LYS and SC-MET media to select haploid spores. This assay enabled subsequent mating of humanized haploids to each other and also generating homozygous diploids for the human genes *via* CRISPR-Cas9. All heterozygous diploids were scored for spore viability and 2:2 segregation of LYS, MET, and MAT loci.

### Gene tree construction

Maximum likelihood trees for each cytoskeletal family were constructed with the Gamma LG protein model with the rapid hill-climbing algorithm bootstrapping 1000 replicates on human and yeast protein sequences curated from the Uniprot database. All trees were computed using Geneious v10.2.6’s RAxML’s plugin^75^ (v8.2.11).

### Human cytoskeleton RNA expression analysis

RNA expression profiles across human tissue types were obtained from the Human Protein Atlas^76^. We extracted the human cytoskeletal gene set from the available expression data normalized via FPKM. The normalized data was then processed in Rstudio and plotted with ggplot2 using the ggridges package.

### SCMD-SGD database searching and evaluations

Genetic interaction data for each assayed yeast cytoskeletal gene was downloaded from the Saccharomyces Genome Database^4, 69^ (SGD) to curate all the interaction partners for the assayed yeast cytoskeletal gene set. Imaging data and morphology parameter files were downloaded from the Saccharomyces Morphological Database^77^ (SCMD) website. As relevant parameters to the phenotypes observed in the case of actin and septin interactors, we specifically examined the long (C103) and short axis (C104) lengths of mother cells budded (SCMD cell types B and C) and unbudded cells (SCMD cell type A). C103/C104 ratios were calculated and ranked in ascending order. We reasoned that deletion strains resulting in small cells with round phenotypes (similar to our humanized actin yeast strains) would have ratios close to 1. In contrast, abnormally shaped cells (similar to our humanized septin yeast strains) would have larger (>1) C103/C104 ratios. We ranked the deletion collection phenotypes by the long to short axis ratios of the budded cells as measure of circularity. Significance was determined by the hypergeometric test (**Table S4**).

## Acknowledgements

The authors would like to thank Charles Boone (University of Toronto) for generously sharing the yeast temperature-sensitive collection and Maitreya Dunham (University of Washington) for insightful discussions and feedback. The authors would also like to thank Angela Bardo, Jagganath Swaminathan, Fan Tu, and the UT ICMB microscopy core for assistance with fluorescence microscopy and imaging. The authors would also like to thank Christopher Yellman (The University of Texas at Austin) for sharing yeast strains in addition to useful discussions. This research was funded by the American Heart Association Predoctoral fellowship (#18PRE34060258) to R.K.G, Natural Sciences and Engineering Research Council (NSERC) of Canada Discovery grant [RGPIN-2018-05089], CRC Tier 2 [NSERC/CRSNG-950-231904] and Canada Foundation for Innovation and Québec Ministère de l’Économie, de la Science et de l’Innovation (#37415) to A.H.K. and from the Welch Foundation (F-1515) and National Institutes of Health (R35 GM122480) to E.M.M.

## Figure legends

**Figure S1.**
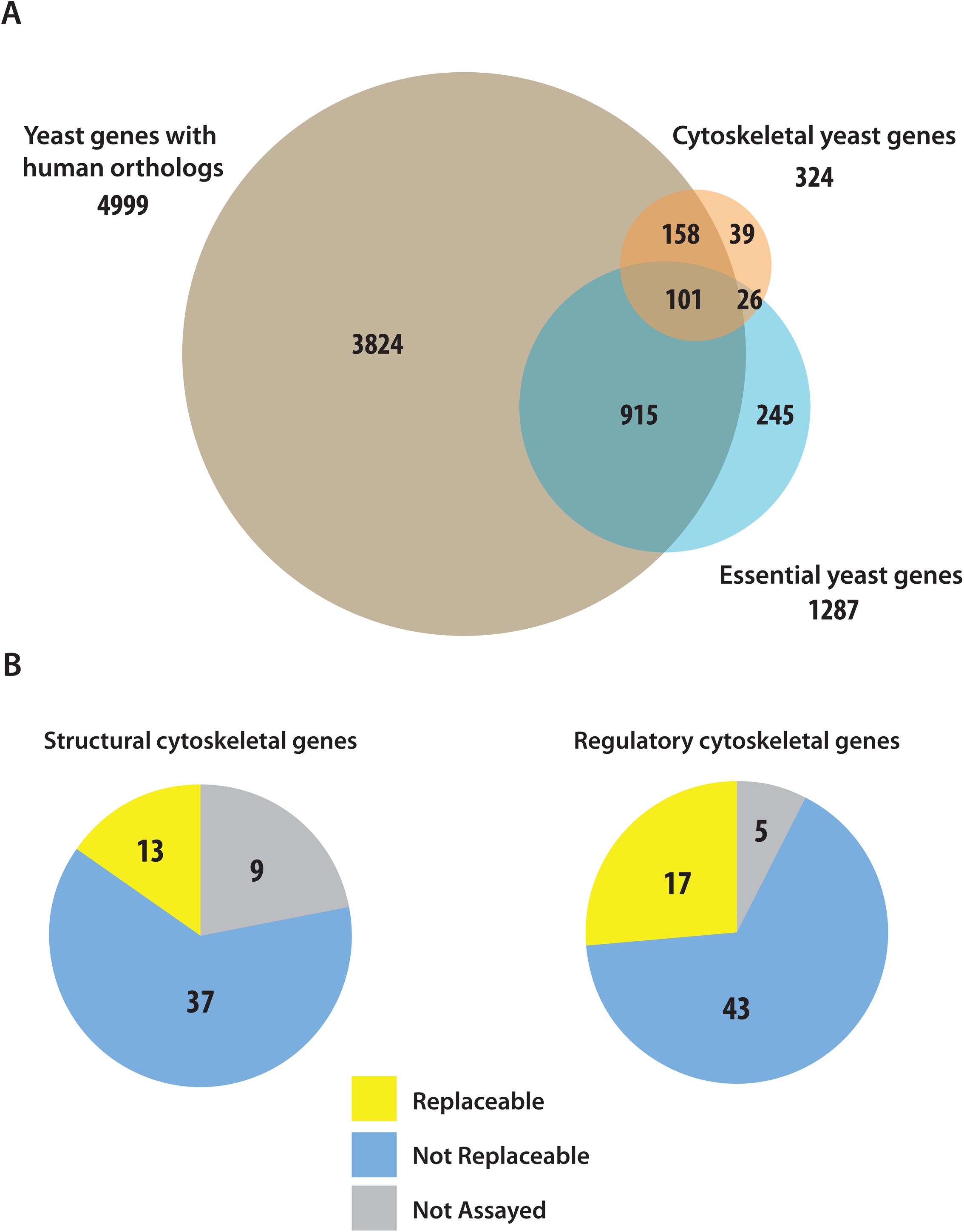
Humans and yeast share a substantial number of orthologous genes. (A) Global orthology relationships shared between human and yeast genes. Yeast genes with human orthologs are represented in brown, cytoskeleton, and essential yeast genes in orange and blue, respectively. (B) Replaceability distribution of human genes across structural and regulatory elements of the eukaryotic cytoskeleton. Data for regulatory human cytoskeletal elements from Laurent *et al*^11^.

**Figure S2.**
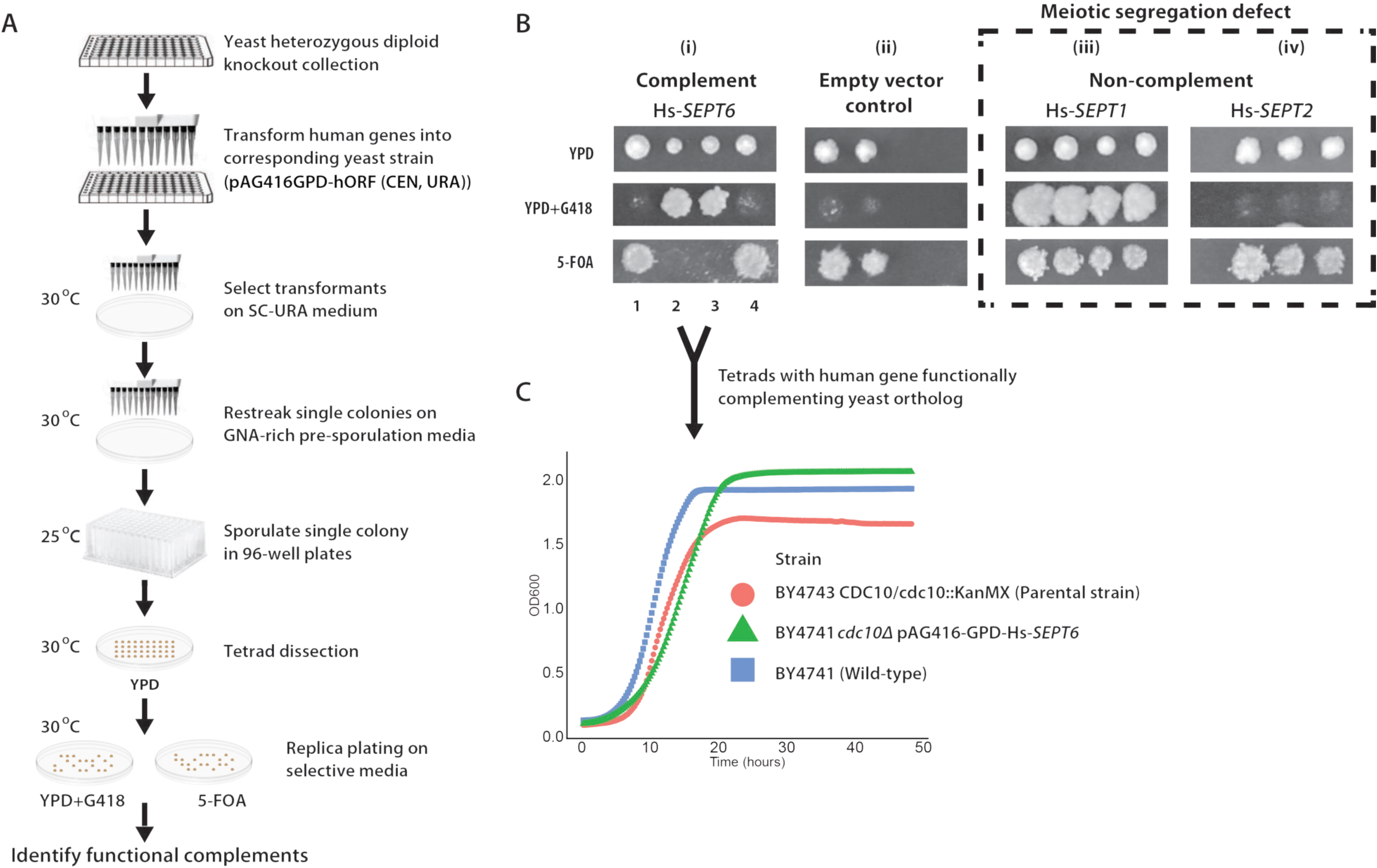
Representative heterozygous diploid deletion allele complementation assay. (A) Schematic outline of humanization assays performed in heterozygous deletion allele (“Magic Marker”) yeast strains. (B) Complementation assay performed after sporulation and tetrad dissection shows examples of (i) replaceable, (ii) empty vector control, and (iii) non-complementing cases. (C) Representative liquid growth assay of humanized strains (Green triangle) compared to the wild-type haploid (blue squares) and parental heterozygous diploid (red circles) yeast strains. Solid shapes represent mean OD600, while shaded boundaries denote +/− standard deviation of 3 replicates.

**Figure S3.**
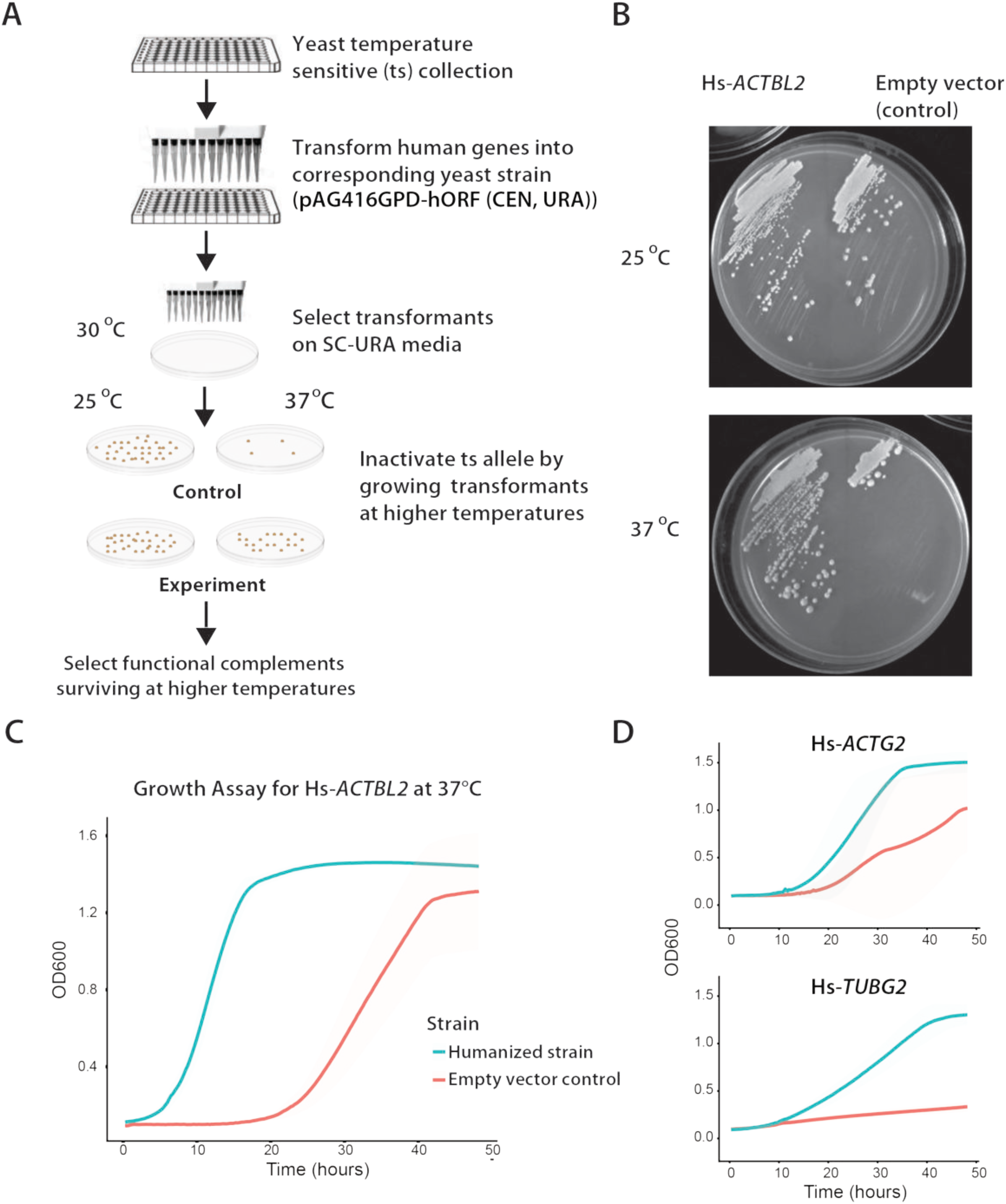
Representative temperature-sensitive allele complementation assay. (A) Schematic outline of the assays performed in temperature-sensitive haploid yeast strains. (B) *Hs-ACTBL2* functionally replaces the temperature dependent lethal growth defect. (C) and (D) Growth assays of humanized strains (blue) and empty vector control (red) at 37°C. Solid lines represent mean OD600, while shaded boundaries denote +/− standard deviation of 3 replicates.

**Figure S4.**
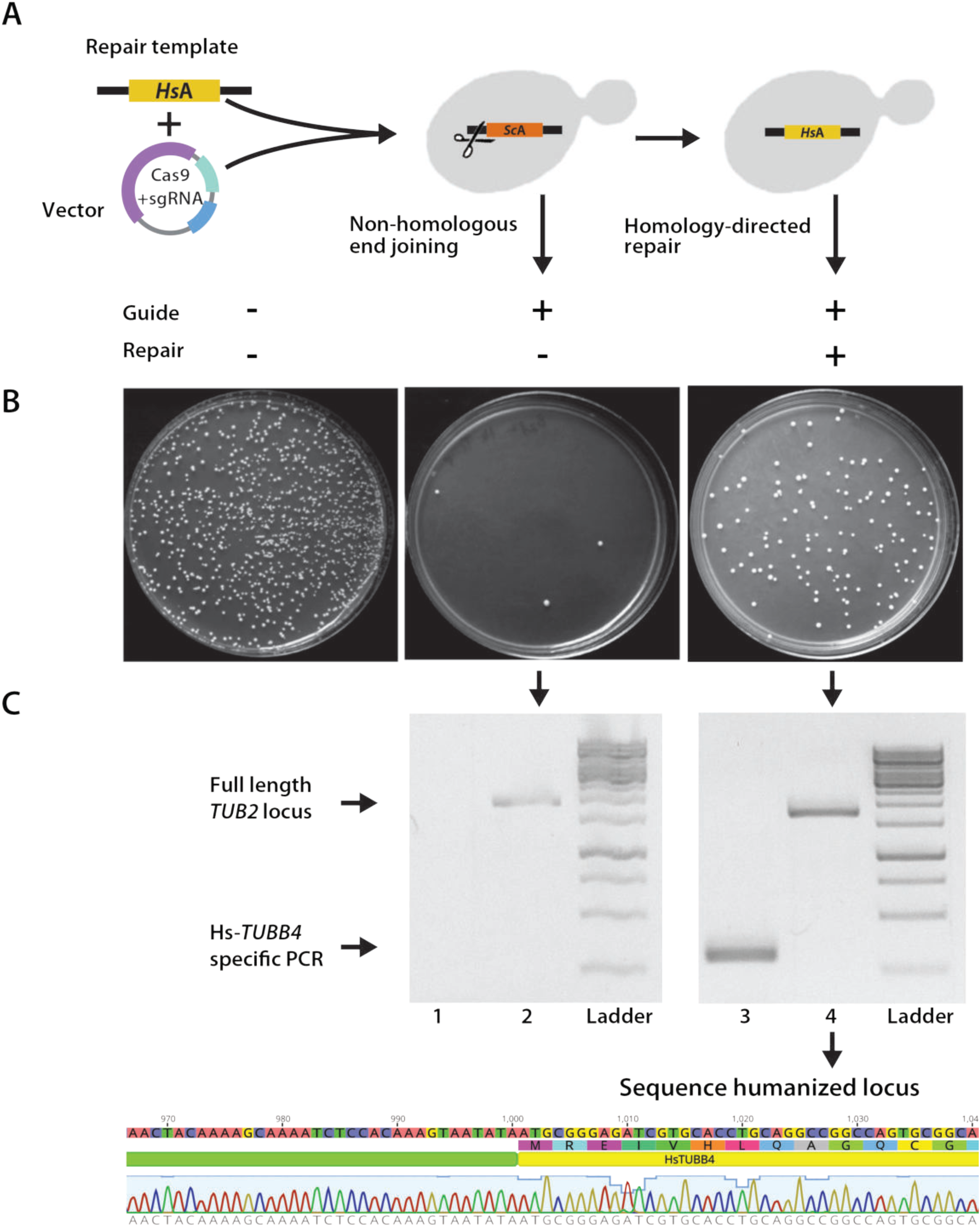
Representative CRISPR-Cas9 complementation assay. (A) Schematic outline of humanization of yeast genes at their native yeast loci using CRISPR-Cas9 based genome editing (B) Yeast cells are rescued from lethal double stranded breaks caused by the Cas9-sgRNA complex (center plate) by supplying a human ortholog containing repair template (right plate) with flanking sequence homology to yeast locus of interest. The left plate is a negative control (carrying the same selectable marker) without Cas9 and sgRNA expressing transcription units, performed to estimate competent cell transformation efficiency. (C) Colony PCR (top) and Sanger sequencing (bottom) to confirm the integration of the human ortholog Hs-*TUBB4* (Lane 3) into the corresponding yeast *TUB2* genomic locus (Lane 4) using primers binding outside the region of homology provided for the repair of the double strand break.

**Figure S5.**
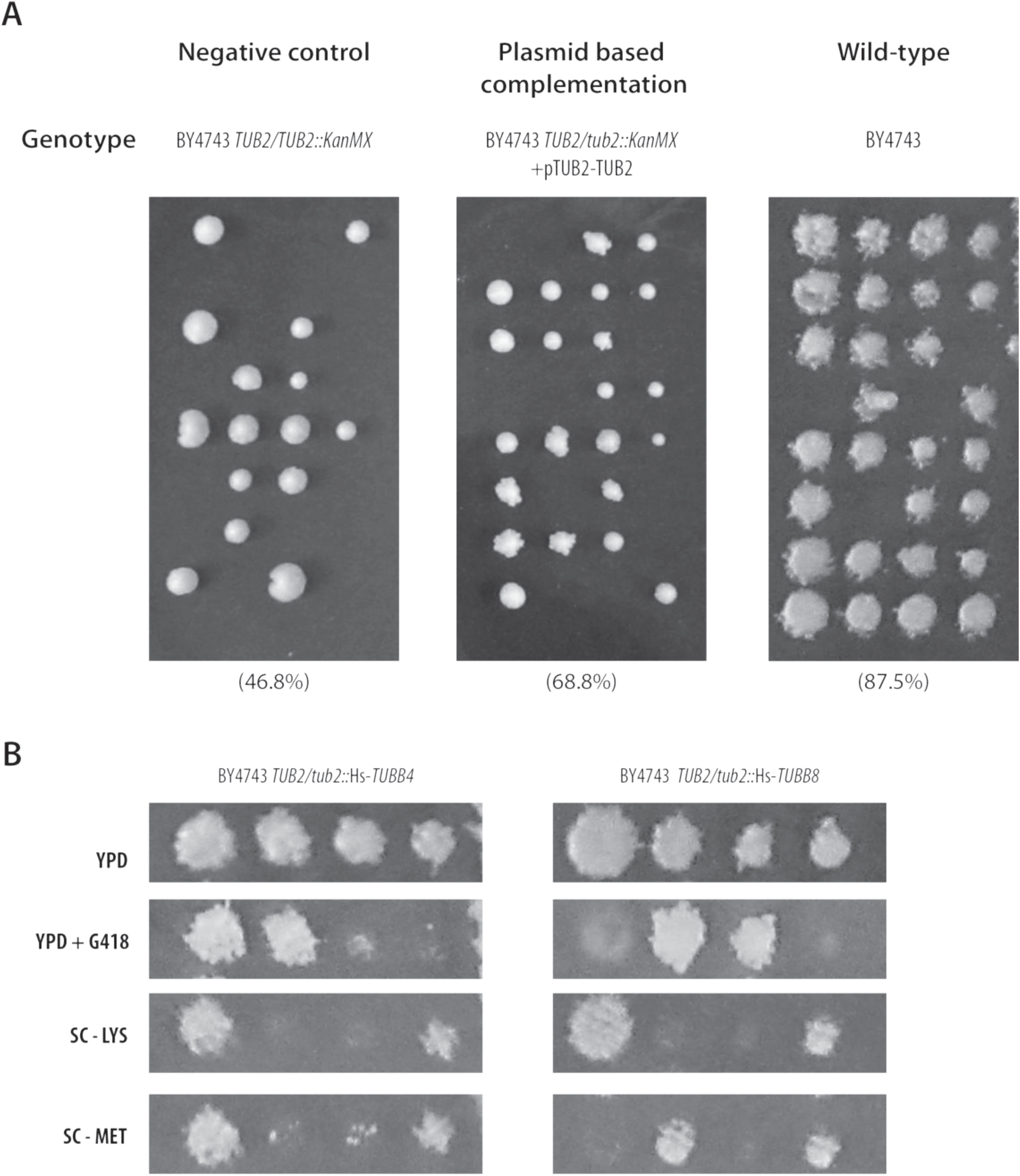
Sporulation and mating efficiencies of humanized tubulin strains. (A) Sporulation and tetrad dissection of BY4743 *TUB2*/*tub2Δ* heterozygous knockout Magic marker strain serves as a negative control in this experiment (left panel) and the yeast strain harboring plasmid borne copy of the yeast *TUB2* gene (center panel). Wild-type homozygous diploid (right panel) serves as a positive control. Plasmid borne *TUB2* only partially rescues the loss-of-function of the yeast gene at the native locus even though it is regulated by the native yeast promoter. (B) Humanized strains heterozygous diploid for Hs-*TUBB4* (left panel) and Hs-*TUBB8* (right panel) were sporulated, tetrad-dissected, and selected on MET^−^ (harbored by BY4741) and LYS^−^ (harbored by BY4742) media to check individual loci for 2:2 independent segregation. Since we performed tubulin humanizations in a BY4741 *tub3*::KanMX background, we also scored G418 resistance, which also segregated 2:2 manner.

**Figure S6.**
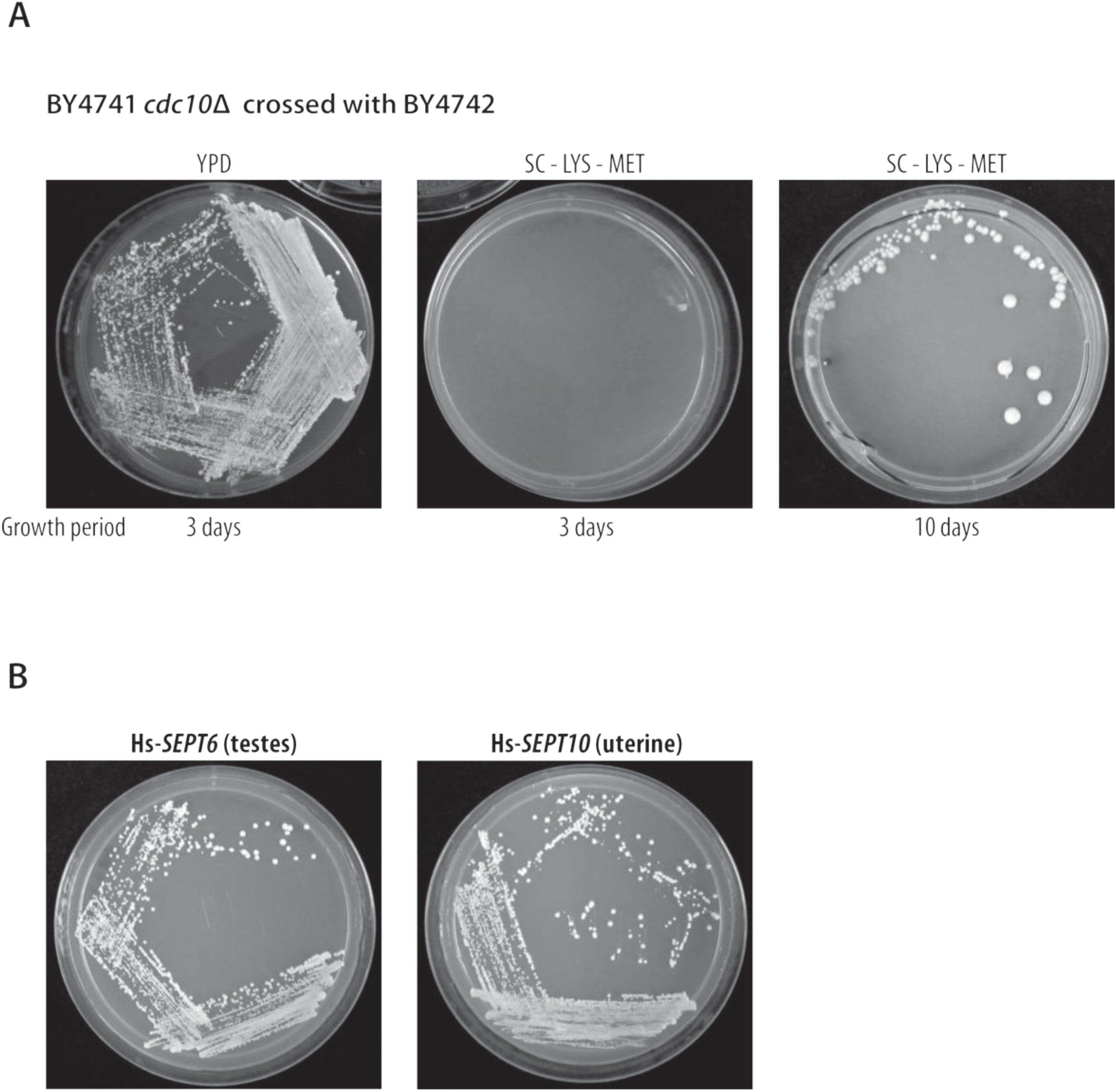
Humanized septin mating assays. (A) Mating assays performed with BY4741 *cdc10Δ* strains show severe mating defects resulting from deletion of the yeast septin *CDC10*. (B) However, human septin isoforms along with other septin, Hs-*SEPT6* and Hs-*SEPT10* can individually rescue the mating defect caused by the deletion of the yeast ortholog, *CDC10*.

**Figure S7.**
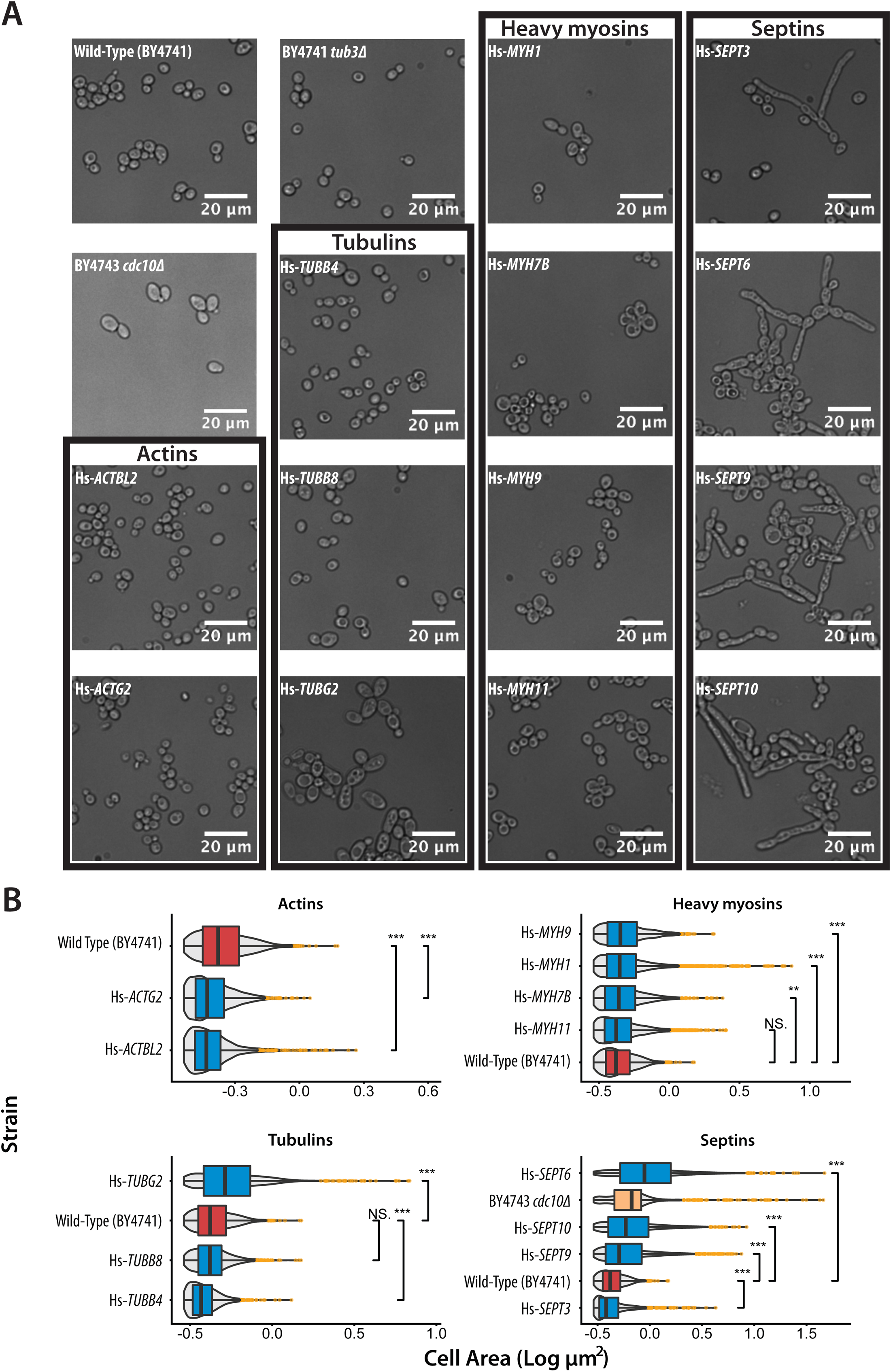
Yeast strains with replaceable human cytoskeletal genes exhibit abnormal cell morphologies. (A) Bright-field images of humanized yeast strains sorted by family. (B) Pairwise comparisons of cell size distributions between humanized and wild-type yeast strains across cytoskeletal families sorted by median size with grey violins indicating cell size. Outliers are indicated by orange dots. Significance comparisons with wild-type were calculated using standard t-test.

**Figure S8.**
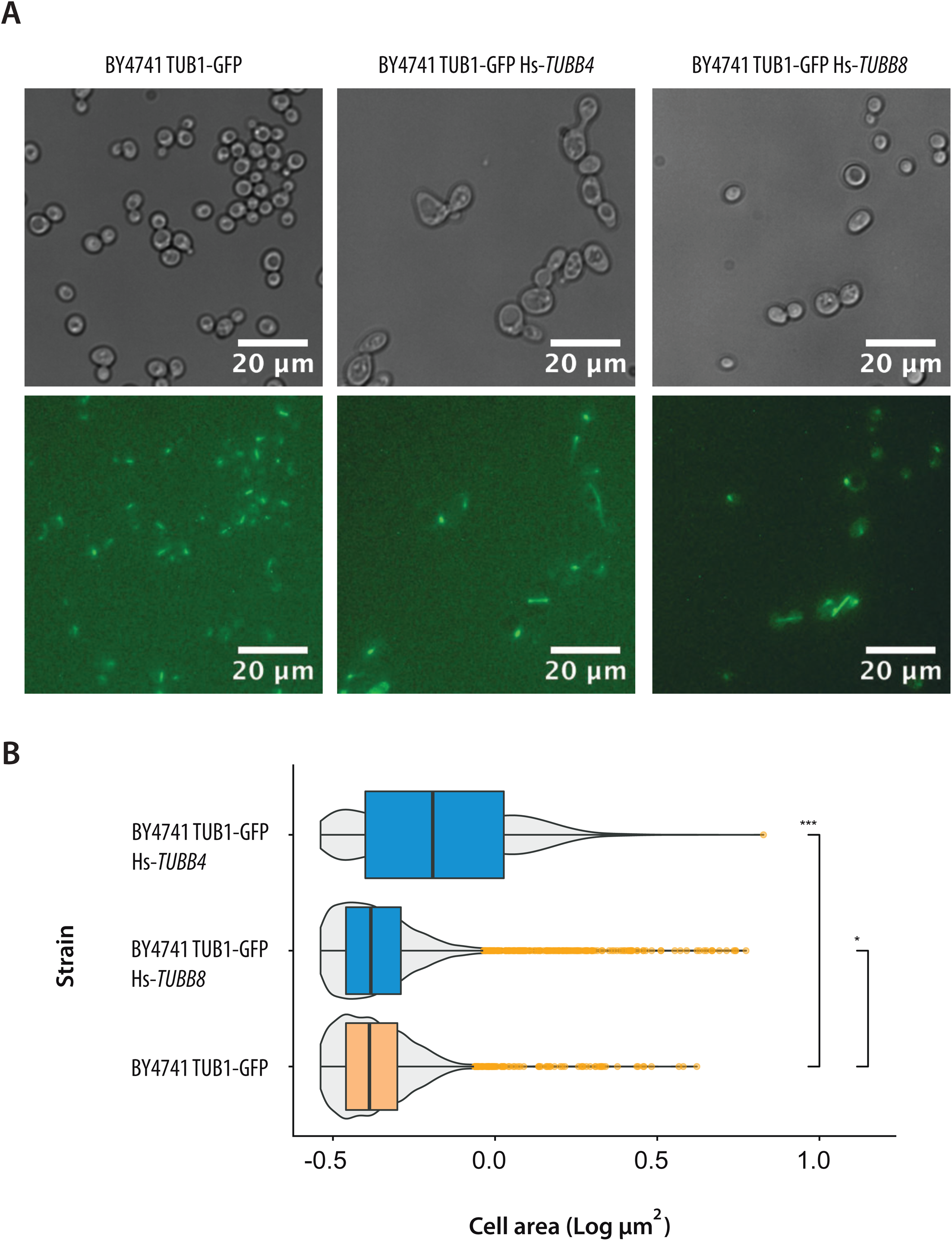
Human β-tubulins are successfully incorporated into the microtubule structure. (A) GFP-fluorescence imaging of the wild-type haploid yeast strain with yeast α-tubulin *TUB1* tagged with GFP (left panel) and harboring human β-tubulins, Hs-*TUBB4* (center panel) and Hs-*TUBB8* (right panel). (B) Cell size distributions of the strains imaged in (A) reveal an obvious cell size increase when replacing Hs-*TUBB4* in a *TUB1-GFP* background. Grey violins indicate size distributions. Significance comparisons were calculated using standard t-test.

**Figure S9.**
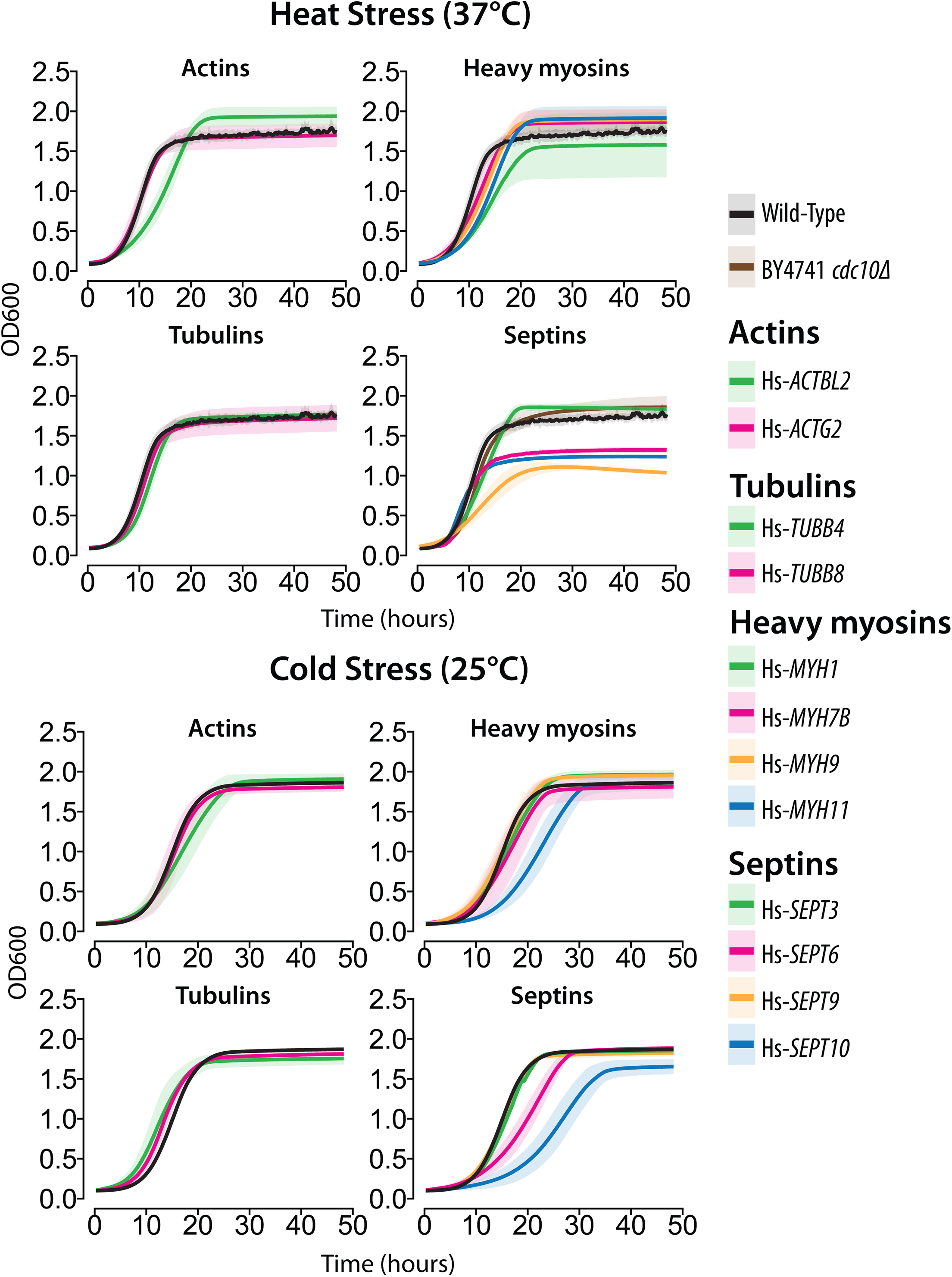
Humanized cytoskeletal strains demonstrate temperature dependent fitness defects. Quantitative growth assays on humanized strains subjected heat (37°C) and cold (25°C) stress. The mean (solid line) +/− standard deviation (shaded boundary) is plotted for 3 replicates.

**Figure S10.**
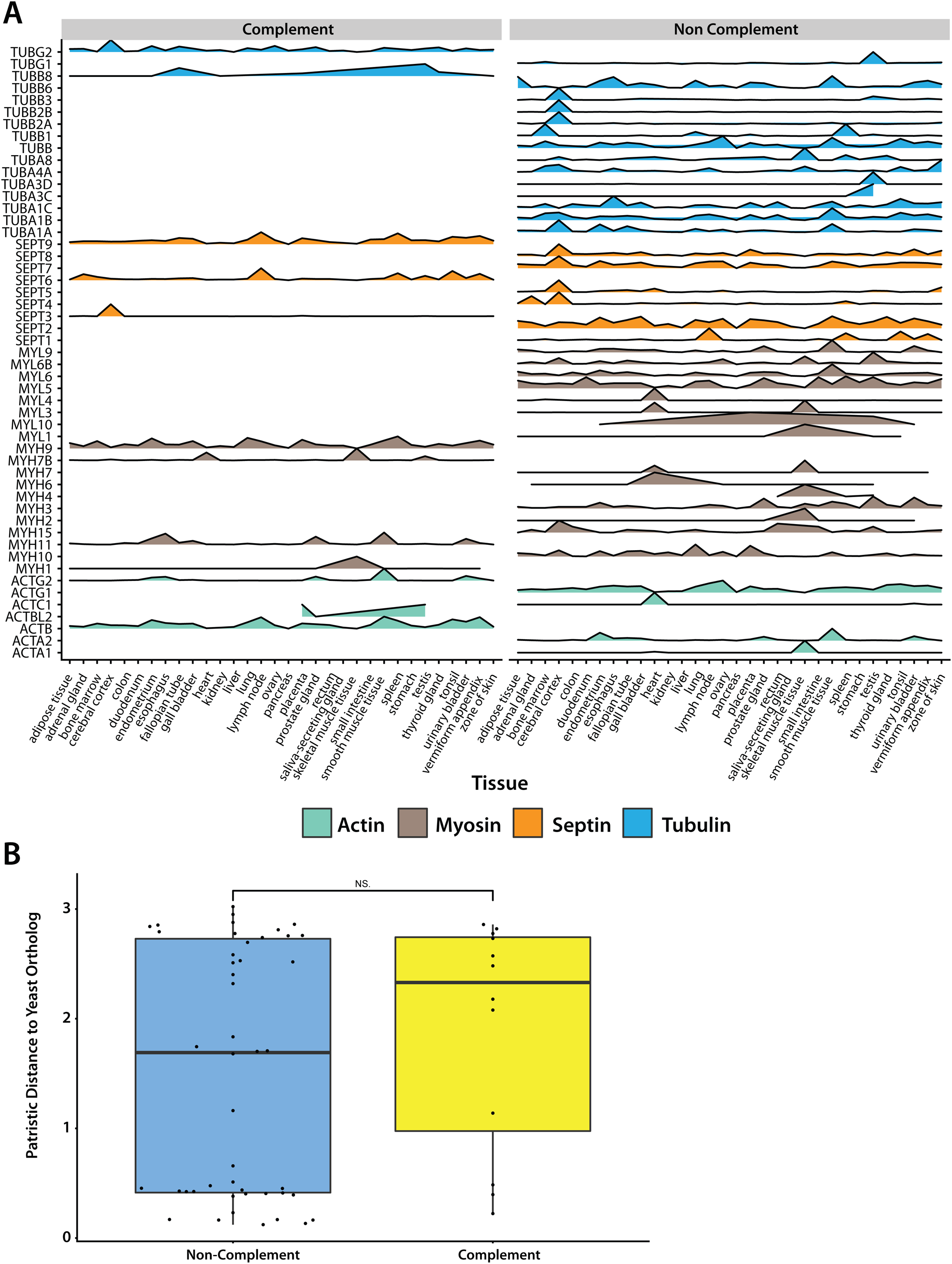
(A)Tissue expression patterns of human cytoskeletal genes do not explain replaceability. Tissue expression data was curated from Human Protein Atlas for available human cytoskeletal genes. X axis represents tissue type, Y axis represents cytoskeletal gene. Ridgeplots are binned by complementation status. (B) Patristic distance between human and yeast ortholog pairs does not predict replaceability. Boxplot comparing patristic distances of a human-yeast ortholog pairs in cytoskeletal families. Patristic distance computed from multiple sequence alignment from MAFFT.

## References

1. Soukup, S. W. Evolution by gene duplication. S. Ohno. Springer-Verlag, New York. 1970. 160 pp. Teratology 9, 250–251.

2. Koonin, E. V. Orthologs, Paralogs, and Evolutionary Genomics. Annu. Rev. Genet. 39, 309–338 (2005).

3. Heinicke, S. et al. The Princeton Protein Orthology Database (P-POD): A Comparative Genomics Analysis Tool for Biologists. PLOS ONE 2, e766 (2007).

4. Cherry, J. M. et al. Saccharomyces Genome Database: the genomics resource of budding yeast. Nucleic Acids Res 40, D700–D705 (2012).

5. Yang, F. et al. Identifying pathogenicity of human variants via paralog-based yeast complementation. PLOS Genetics 13, e1006779 (2017).

6. Sun, S. et al. An extended set of yeast-based functional assays accurately identifies human disease mutations. Genome Res. 26, 670–680 (2016).

7. Hamza, A. et al. Complementation of Yeast Genes with Human Genes as an Experimental Platform for Functional Testing of Human Genetic Variants. Genetics 201, 1263–1274 (2015).

8. Kachroo, A. H. et al. Systematic bacterialization of yeast genes identifies a near-universally swappable pathway. eLife 6, e25093 (2017).

9. Kachroo, A. H. et al. Systematic humanization of yeast genes reveals conserved functions and genetic modularity. Science 348, 921–925 (2015).

10. Laurent, J. M., Young, J. H., Kachroo, A. H. & Marcotte, E. M. Efforts to make and apply humanized yeast. Brief Funct Genomics 15, 155–163 (2016).

11. Laurent, J. M. et al. Humanization of yeast genes with multiple human orthologs reveals principles of functional divergence between paralogs. bioRxiv 668335 (2019) doi:10.1101/668335.

12. Wickstead, B. & Gull, K. The evolution of the cytoskeleton. The Journal of Cell Biology 194, 513–525 (2011).

13. Wickstead, B., Gull, K. & Richards, T. A. Patterns of kinesin evolution reveal a complex ancestral eukaryote with a multifunctional cytoskeleton. BMC Evolutionary Biology 10, 110 (2010).

14. Wickstead, B. & Gull, K. Dyneins Across Eukaryotes: A Comparative Genomic Analysis. Traffic 8, 1708–1721.

15. Jékely, G. Origin and Evolution of the Self-Organizing Cytoskeleton in the Network of Eukaryotic Organelles. Cold Spring Harb Perspect Biol 6, a016030 (2014).

16. Hall, P. A. & Russell, S. H. The pathobiology of the septin gene family. The Journal of Pathology 204, 489–505.

17. McKean, P. G., Vaughan, S. & Gull, K. The extended tubulin superfamily. Journal of Cell Science 114, 2723–2733 (2001).

18. Janke, C. The tubulin code: Molecular components, readout mechanisms, and functions. J Cell Biol 206, 461–472 (2014).

19. Winsor, B. & Schiebel, E. Review: An Overview of the Saccharomyces cerevisiae Microtubule and Microfilament Cytoskeleton. Yeast 13, 399–434 (1997).

20. Schatz, P. J., Pillus, L., Grisafi, P., Solomon, F. & Botstein, D. Two functional alpha-tubulin genes of the yeast Saccharomyces cerevisiae encode divergent proteins. Mol. Cell. Biol. 6, 3711–3721 (1986).

21. Bode, C. J., Gupta, M. L., Suprenant, K. A. & Himes, R. H. The two α-tubulin isotypes in budding yeast have opposing effects on microtubule dynamics in vitro. EMBO reports 4, 94–99 (2003).

22. Li, Z. et al. Systematic exploration of essential yeast gene function with temperature-sensitive mutants. Nat Biotech 29, 361–367 (2011).

23. Nogales, E., Ramey, V. H. & Wang, H.-W. Chapter 8 - Cryo-EM Studies of Microtubule Structural Intermediates and Kinetochore–Microtubule Interactions. in Methods in Cell Biology (ed. Correia, L. W. and J. J.) vol. 95 128–156 (Academic Press, 2010).

24. Chen, B. et al. The comprehensive mutational and phenotypic spectrum of TUBB8 in female infertility. European Journal of Human Genetics 27, 300 (2019).

25. Feng, R. et al. Mutations in TUBB8 and Human Oocyte Meiotic Arrest. New England Journal of Medicine 374, 223–232 (2016).

26. Yuan, P. et al. A novel mutation in the TUBB8 gene is associated with complete cleavage failure in fertilized eggs. J Assist Reprod Genet 35, 1349–1356 (2018).

27. Stottmann, R. W. et al. A Heterozygous Mutation in Tubulin, Beta 2b (tubb2b) Causes Cognitive Deficits and Hippocampal Disorganization. Genes, Brain and Behavior n/a-n/a (2016) doi:10.1111/gbb.12327.

28. Huang, H. et al. Human TUBB3 Mutations Disrupt Netrin Attractive Signaling. Neuroscience 374, 155–171 (2018).

29. Tischfield, M. A. et al. Human TUBB3 Mutations Perturb Microtubule Dynamics, Kinesin Interactions, and Axon Guidance. Cell 140, 74–87 (2010).

30. Wawro, M. E. et al. Tubulin beta 3 and 4 are involved in the generation of early fibrotic stages. Cellular Signalling 38, 26–38 (2017).

31. Huerta-Cepas, J. et al. eggNOG 4.5: a hierarchical orthology framework with improved functional annotations for eukaryotic, prokaryotic and viral sequences. Nucleic Acids Res 44, D286–D293 (2016).

32. Sonnhammer, E. L. L. & Östlund, G. InParanoid 8: orthology analysis between 273 proteomes, mostly eukaryotic. Nucleic Acids Res 43, D234–D239 (2015).

33. Giaever, G. et al. Functional profiling of the Saccharomyces cerevisiae genome. Nature 418, 387–391 (2002).

34. Winzeler, E. A. et al. Functional Characterization of the S. cerevisiae Genome by Gene Deletion and Parallel Analysis. Science 285, 901–906 (1999).

35. Wach, A., Brachat, A., Pöhlmann, R. & Philippsen, P. New heterologous modules for classical or PCR-based gene disruptions in Saccharomyces cerevisiae. Yeast 10, 1793–1808 (1994).

36. Kofoed, M. et al. An Updated Collection of Sequence Barcoded Temperature-Sensitive Alleles of Yeast Essential Genes. G3 5, 1879–1887 (2015).

37. Akhmetov, A. et al. Single-step Precision Genome Editing in Yeast Using CRISPR-Cas9. BIO-PROTOCOL 8, (2018).

38. Lamesch, P. et al. hORFeome v3.1: A resource of human open reading frames representing over 10,000 human genes. Genomics 89, 307–315 (2007).

39. Rual, J.-F. et al. Towards a proteome-scale map of the human protein–protein interaction network. Nature 437, 1173–1178 (2005).

40. Team, T. M. P. et al. The completion of the Mammalian Gene Collection (MGC). Genome Res. 19, 2324–2333 (2009).

41. Neff, N. F., Thomas, J. H., Grisafi, P. & Botstein, D. Isolation of the β-tubulin gene from yeast and demonstration of its essential function in vivo. Cell 33, 211–219 (1983).

42. Vogel, J. et al. Phosphorylation of γ-Tubulin Regulates Microtubule Organization in Budding Yeast. Developmental Cell 1, 621–631 (2001).

43. Bardwell, L. A walk-through of the yeast mating pheromone response pathway. Peptides 26, 339–350 (2005).

44. Molk, J. N. & Bloom, K. Microtubule dynamics in the budding yeast mating pathway. J Cell Sci 119, 3485–3490 (2006).

45. Neiman, A. M. Sporulation in the Budding Yeast Saccharomyces cerevisiae. Genetics 189, 737–765 (2011).

46. Pan, F., Malmberg, R. L. & Momany, M. Analysis of septins across kingdoms reveals orthology and new motifs. BMC Evol Biol 7, 103 (2007).

47. McMurray, M. A. et al. Septin Filament Formation Is Essential in Budding Yeast. Developmental Cell 20, 540–549 (2011).

48. Douglas, L. M., Alvarez, F. J., McCreary, C. & Konopka, J. B. Septin Function in Yeast Model Systems and Pathogenic Fungi. Eukaryotic Cell 4, 1503–1512 (2005).

49. Kim, J. & Rose, M. D. Stable Pseudohyphal Growth in Budding Yeast Induced by Synergism between Septin Defects and Altered MAP-kinase Signaling. PLOS Genetics 11, e1005684 (2015).

50. McMurray, M. A. & Thorner, J. Septin Stability and Recycling during Dynamic Structural Transitions in Cell Division and Development. Current Biology 18, 1203–1208 (2008).

51. Weems, A. & McMurray, M. The step-wise pathway of septin hetero-octamer assembly in budding yeast. eLife 6, e23689 (2017).

52. Huh, W.-K. et al. Global analysis of protein localization in budding yeast. Nature 425, 686–691 (2003).

53. Caudron, F. et al. Mutation of Ser172 in yeast β tubulin induces defects in microtubule dynamics and cell division. PLoS ONE 5, e13553 (2010).

54. Kiss, E. et al. Comparative genomics reveals the origin of fungal hyphae and multicellularity. Nature Communications 10, 1–13 (2019).

55. Mukaremera, L., Lee, K. K., Mora-Montes, H. M. & Gow, N. A. R. Candida albicans Yeast, Pseudohyphal, and Hyphal Morphogenesis Differentially Affects Immune Recognition. Front. Immunol. 8, (2017).

56. Warenda, A. J. & Konopka, J. B. Septin Function in Candida albicans Morphogenesis. Mol Biol Cell 13, 2732–2746 (2002).

57. Beilharz, T. H. et al. Coordination of Cell Cycle Progression and Mitotic Spindle Assembly Involves Histone H3 Lysine 4 Methylation by Set1/COMPASS. Genetics 205, 185–199 (2017).

58. Burke, D., Gasdaska, P. & Hartwell, L. Dominant effects of tubulin overexpression in Saccharomyces cerevisiae. Mol Cell Biol 9, 1049–1059 (1989).

59. Weinstein, B. & Solomon, F. Phenotypic consequences of tubulin overproduction in Saccharomyces cerevisiae: differences between alpha-tubulin and beta-tubulin. Molecular and Cellular Biology 10, 5295–5304 (1990).

60. Abruzzi, K. C., Smith, A., Chen, W. & Solomon, F. Protection from Free β-Tubulin by the β-Tubulin Binding Protein Rbl2p. Mol Cell Biol 22, 138–147 (2002).

61. Kaplan, C., Steinmann, M., Zapiorkowska, N. A. & Ewers, H. Functional Redundancy of Septin Homologs in Dendritic Branching. Front. Cell Dev. Biol. 5, (2017).

62. Fujishima, K., Kiyonari, H., Kurisu, J., Hirano, T. & Kengaku, M. Targeted disruption of Sept3, a heteromeric assembly partner of Sept5 and Sept7 in axons, has no effect on developing CNS neurons. Journal of Neurochemistry 102, 77–92 (2007).

63. Tsang, C. W. et al. Superfluous Role of Mammalian Septins 3 and 5 in Neuronal Development and Synaptic Transmission. Molecular and Cellular Biology 28, 7012–7029 (2008).

64. Falk, J., Boubakar, L. & Castellani, V. Septin functions during neuro-development, a yeast perspective. Current Opinion in Neurobiology 57, 102–109 (2019).

65. Soifer, I. & Barkai, N. Systematic identification of cell size regulators in budding yeast. Mol Syst Biol 10, (2014).

66. Lindsey, R., Ha, Y. & Momany, M. A Septin from the Filamentous Fungus A. nidulans Induces Atypical Pseudohyphae in the Budding Yeast S. cerevisiae. PLOS ONE 5, e9858 (2010).

67. Sun, S. et al. A proactive genotype-to-patient-phenotype map for cystathionine beta-synthase. bioRxiv 473983 (2018) doi:10.1101/473983.

68. Marzo, M. G. et al. Molecular basis for dyneinopathies reveals insight into dynein regulation and dysfunction. eLife 8, e47246 (2019).

69. Cherry, J. M. et al. SGD: Saccharomyces Genome Database. Nucleic Acids Res 26, 73–79 (1998).

70. Alberti, S., Gitler, A. D. & Lindquist, S. A suite of Gateway® cloning vectors for high-throughput genetic analysis in Saccharomyces cerevisiae. Yeast 24, 913–919 (2007).

71. Methods in Yeast Genetics and Genomics: A Cold Spring Harbor Laboratory Course Manual, 2015 Edition. (Cold Spring Harbor Laboratory Press, 2015).

72. Lee, M. E., DeLoache, W. C., Cervantes, B. & Dueber, J. E. A Highly Characterized Yeast Toolkit for Modular, Multipart Assembly. ACS Synth. Biol. 4, 975–986 (2015).

73. Schindelin, J., Rueden, C. T., Hiner, M. C. & Eliceiri, K. W. The ImageJ ecosystem: An open platform for biomedical image analysis. Molecular Reproduction and Development 82, 518–529 (2015).

74. Schindelin, J. et al. Fiji: an open-source platform for biological-image analysis. Nature Methods 9, 676–682 (2012).

75. Stamatakis, A. RAxML version 8: a tool for phylogenetic analysis and post-analysis of large phylogenies. Bioinformatics 30, 1312–1313 (2014).

76. Uhlén, M. et al. Tissue-based map of the human proteome. Science 347, 1260419 (2015).

77. Saito, T. L. et al. SCMD: Saccharomyces cerevisiae Morphological Database. Nucleic Acids Res. 32, D319–322 (2004).

78. Agapite, J. et al. Alliance of Genome Resources Portal: unified model organism research platform. Nucleic Acids Res doi:10.1093/nar/gkz813.

